# Assembly of a persistent apical actin network by the formin Frl/Fmnl tunes epithelial cell deformability

**DOI:** 10.1101/680033

**Authors:** Benoit Dehapiot, Raphaël Clément, Gabriella Gazsó-Gerhát, Jean-Marc Philippe, Thomas Lecuit

## Abstract

Tissue remodeling during embryogenesis is driven by the apical contractility of the epithelial cell cortex. This behavior arises notably from Rho1/Rok induced transient accumulation of non-muscle myosin II (MyoII pulses) pulling on actin filaments (F-Actin) of the medio-apical cortex. While recent studies begin to highlight the mechanisms governing the emergence of Rho1/Rok/MyoII pulsatility in different organisms, little is known about how the F-Actin organization influences this process. Focusing on *Drosophila* ectodermal cells during germband extension and amnioserosa cells during dorsal closure, we show that the medio-apical actomyosin cortex consists of two entangled F-Actin subpopulations. One exhibits pulsatile dynamics of actin polymerization in a Rho1 dependent manner. The other forms a persistent and homogeneous network independent of Rho1. We identify the Frl/Fmnl formin as a critical nucleator of the persistent network since modulating its level, in mutants or by overexpression, decreases or increases the network density. Absence of this network yields sparse connectivity affecting the homogeneous force transmission to the cell boundaries. This reduces the propagation range of contractile forces and results in tissue scale morphogenetic defects. Our work sheds new lights on how the F-Actin cortex offers multiple levels of regulation to affect epithelial cells dynamics.

## Introduction

Animal cells can actively modify their shape in order to complete complex processes such as cell migration, division or cell shape changes during tissue morphogenesis. These behaviors arise from the contractile properties of the actomyosin cortex and its ability to build up tension by sliding MyoII molecular motors over anti-parallel arrays of crosslinked actin filaments^1, 2^.

The recent advances in live imaging have shown that cortical contractility can occur in a pulsatile manner, by taking the form of local and transient accumulations of MyoII, known as MyoII pulses. This phenomenon was first described in the *C. elegans* zygote and has since been reported in many other species, in both embryonic and extra-embryonic tissues^3–11^. MyoII pulses can underly a variety of morphogenetic processes, ranging from single cell polarization to tissue scale remodeling. Although recent evidence suggests that MyoII pulses can emerge spontaneously from stochastic fluctuations and local amplification^12–15^, the spatio-temporal pattern of cortical contractility must be controlled in order to produce reproducible morphogenetic outcomes. In most studied systems, this control is achieved through the conserved RhoA GTPase signaling, which activate MyoII via Rho-associated kinase (ROCK) dependent phosphorylations of its regulatory light chain (MyoII-RLC)^1, 11, 13, 15, 16^.

Besides MyoII regulation, another key parameter influencing cortical contractility resides in actin filament network organization and dynamics. Typically, the cortex assembles as a thin network of actin filaments bound to the plasma membrane. The cortical network is both highly plastic and mechanically rigid and confer to the cells the ability to adapt and exert forces on their surrounding environment^2, 17–19^. These remarkable properties stem from the action of more than a hundred actin binding proteins (ABPs) regulating the organization and the turnover of the network’s components. In brief, actin nucleators, such as the Arp2/3 complex or the formin protein family, first promote the polymerization of filaments and can lead, depending on how they operate, to the assembly of networks harboring different levels of ramification (e.g. highly branched for Arp2/3 and sparse for the formins). After being assembled, the network organization can be remodeled by actin bundlers (Fascin, Plastin) or cross-linkers (Filamin, α-Actinin) and its filament turnover regulated by factors like Profilin, capping proteins or ADF/Cofilin^17–19^. Past experimental and theoretical studies have shown that modulating the dynamic organization of F-Actin networks through ABPs, can significantly modify how the MyoII contractility gives rise to cortical tension^2, 20–22^.

In embryonic *Drosophila* epithelial cells, the MyoII pulses appear in the medio-apical (also referred as medial) part of the cell and produce sustained apical constrictions or repeated cycles of apical contraction/relaxation. These two modalities of MyoII pulsatility, together with adherens junctions (AJs) remodeling, give rise to a variety of morphogenetic events such as mesoderm/endoderm invagination, convergent extension or tissue dorsal closure^4, 6, 7^. While the mechanisms underlying the emergence of MyoII pulsatility have been widely studied, little is known however about how the medio-apical F-Actin supports the pulsatile cell contractility and allows spatial transmission of mechanical stresses. In mechanical terms, it has been shown that actin filaments transmit cortical tension over length scales that depend on viscoelastic properties of the cortex^23^. These properties emerge from the spatial organization and the temporal remodeling of the F-Actin networks which are regulated by ABPs. It has also been shown that the cortical F-Actin can affect the level of MyoII activation by serving as a scaffold for the motor-driven advection of regulators such as Rho1 and Rok^13^ or for the recruitment of Rho1 inhibitors, such as RhoGAPs, required for pulse disassembly and pulsation^15^. Here, focusing on two highly pulsatile tissues, namely the ectodermal cells during germband extension (GBE) and amnioserosa cells during dorsal closure (DC), we first the regulation of the medio-apical F-Actin. We next sought to understand how the medial F-Actin network affects MyoII contractile forces within cells, and it supports the propagation of contractile tension to the surrounding tissue.

## Results

### Spatio-temporal dynamics of medio-apical F-Actin

To monitor F-Actin dynamics, we stably expressed the actin binding domain of Utrophin fused to eGFP (eGFP::UtrCH) in living *Drosophila* embryos. This probe offers a good signal-to-noise ratio to observe isolated actin filaments and does not produce abnormal structures when comparing to phalloidin staining (Supplementary Fig. 1a)^24^. In ectodermal and amnioserosa cells, the medio-apical F-Actin forms a network of filaments that can be observed directly under the apical surface of the cell (Fig. 1a-c). In both tissues, the apical cortex was highly dynamic, displaying contractile foci of F-Actin (Supplementary Movies 1,2) and a constant turnover of its filaments (see single filament assembly and disassembly events in Fig. 1d and Supplementary Movies 3).

**Fig. 1.**
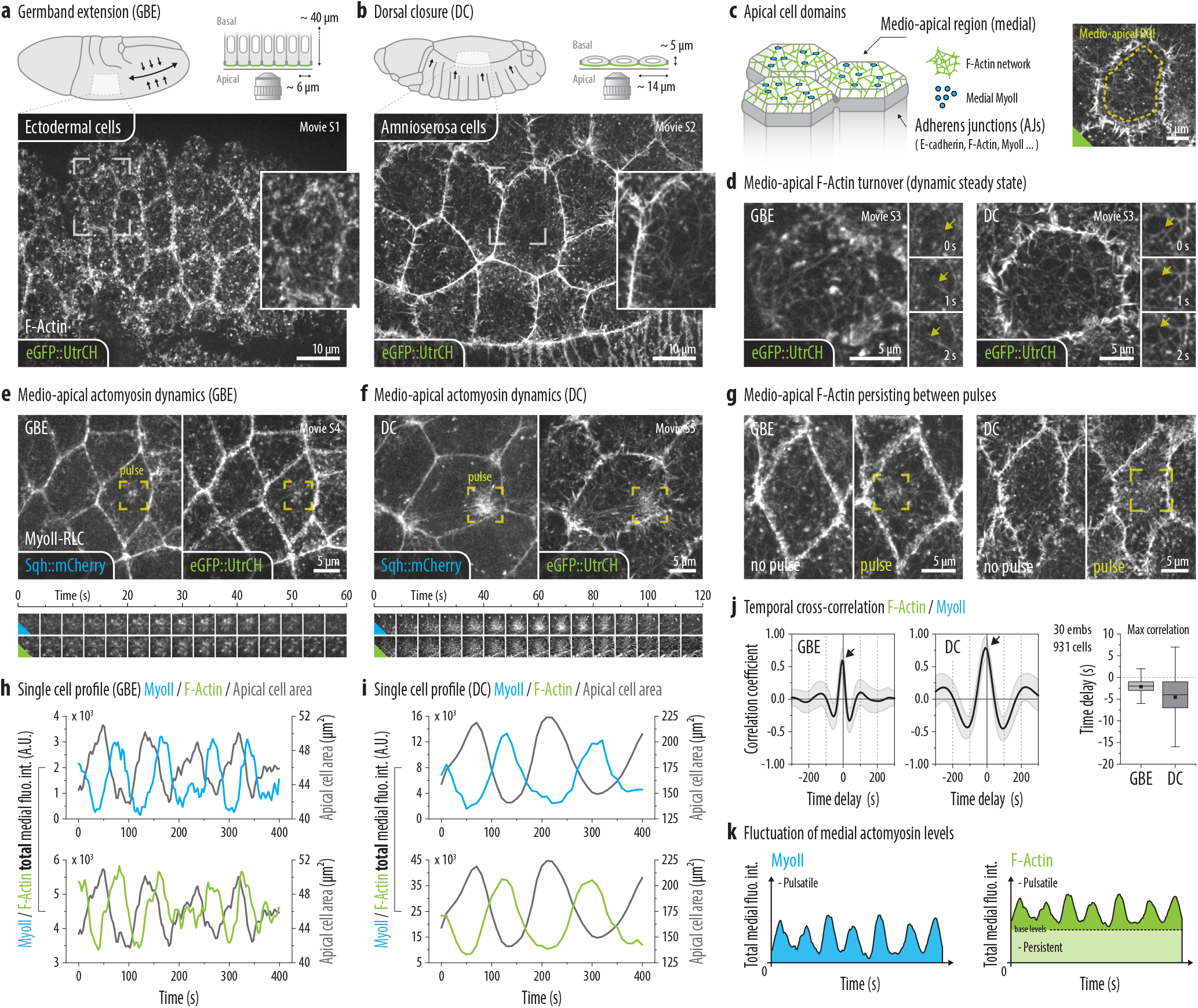
Spatio-temporal dynamics of medio-apical F-Actin. **(a,b)** Live F-Actin localization revealed by the eGFP::UtrCH probe in ectodermal cells during germband elongation (GBE) or amnioserosa cells during dorsal closure (DC). Top panel : schematic of cells localization within the embryo and cross-section of the corresponding epithelia. Main image : single time-point extracted from the Supplementary Movie 1 (GBE) or 2 (DC), showing a max-proj. (4 x 0.33 µm) of the most apical planes. Inserted images : selected zoomed region (see yellow frames, main images). **(c)** Left panel : diagram of the apical domains of early *Drosophila* epithelial cells, representing both the medio-apical and the junctional sub-domains. Right panel : typical region of interest (ROI) used to quantify the medio-apical actomyosin levels. **(d)** High frequency imaging reveals the constant turnover of medio-apical actin filaments (eGFP::UtrCH). Main images : single time-point extracted from the Supplementary Movie 3, left (GBE) or 4 (DC), max-proj. (2 x 0.33 µm). Time series (right) : three consecutive time-points highlighting an actin polymerization event (see white arrows). **(e,f)** Live MyoII-RLC (Sqh::mCherry) and F-Actin (eGFP::UtrCH) localization in cell(s) undergoing cortical pulsed contractility (MyoII pulses). Main images : single time-point extracted from the Supplementary Movie 5 (GBE) or 6 (DC), max-proj. (4 x 0.33 µm). Time series (bottom) : zoomed image sequence showing the assembly/disassembly of a selected actomyosin pulse (see yellow frames, main images). **(g)** Medio-apical F-Actin localization in cell undergoing (pulse) or not (no pulse) a contractile event during GBE (left panel) or DC (right panel). The yellow frames indicate pulses localization. **(h,i)** Temporal variations of apical cell area and total medial MyoII-RLC (top) or F-Actin (bottom) fluo. int. in a selected cell during GBE (g) or DC (h). **(j)** Line graphs : mean ± S.D. of averaged temporal cross-correlation analysis between total medial F-Actin and MyoII-RLC fluo. int. Box plot : time delay for max. correlation of the averaged temporal cross-correlation (see black arrow). **(k)** Schematic representation of the medial actomyosin levels fluctuation. Both MyoII and F-Actin are pulsatile but, contrary to MyoII, the F-Actin oscillates over nonzero baseline of persistent actin filaments. Box plots (i,j) : extend from 1^st^ (Q1) to 3^rd^ (Q3) quartile (Q3-Q1 = IQR), whiskers : Q1 or Q3 ± 1.5 x IQR, horizontal lines : medians, black squares : means.

To understand how the cortex dynamics is influenced by the MyoII pulsed cortical contractility, we co-expressed the utrophin probe with a tagged version of the *Drosophila* MyoII-RLC (Sqh::mCherry or Sqh::mKate2) (Fig. 1e,f and Supplementary Movies 4,5). We first noticed that, contrary to MyoII, the medio-apical F-Actin network persists between cycles of apical constriction (see “no pulse” vs “pulse” in Fig. 1g) and still assembles in the rare non-contractile cells. While this observation suggests a decoupling between the assembly of the medial F-Actin and the emergence of pulsed contractility, we also noticed that the contractile foci of F-Actin correlate with the appearance of MyoII pulses (see yellow frames Fig. 1e,f). Although this could stem from by the tendency of MyoII to advect material while contracting, the amount of F-Actin contained in these foci seems to exceed what one would expect from the simple concentration of molecules. Thus, to further investigate this phenomenon, we designed an automated cell segmentation and background subtraction procedure to carefully measure the actomyosin levels in the restricted medio-apical domain (see Methods and Supplementary Fig. 1b,c). By comparing single cell intensity profiles (Fig. 1h,i) or by performing cross-correlation analysis (Fig. 1j), we found that the F-Actin and the MyoII levels are strongly correlated in time and peak together, with a maximum shift of a few seconds. Since our measurements were carried out on total integrated fluorescence intensities, this observation suggests that a surge of actin polymerization accompanies the formation of MyoII pulses. Overall, we concluded that medio-apical F-Actin exhibits two distinct behaviors. First, cells assemble a persistent network of actin filaments, in the form of a homogeneous network. Second, cells induce pulsatile F-actin polymerization in synchrony with MyoII pulses (see diagram in Fig. 1k).

### Rho1 pathway inhibition reveals two differentially regulated medio-apical F-Actin sub-populations

To further characterize the mechanisms underlying medio-apical F-Actin dynamics, we inhibited MyoII pulsatility by targeting molecular components of the Rho1 signaling pathway (Fig. 2a). We tested whether the pulsatile accumulation of F-Actin and MyoII is co-regulated with or decoupled from the mechanisms promoting the assembly of the persistent network. We first focused on ectodermal cells during GBE and designed two different strategies to inhibit Rho1. In the first case, we injected the C3-transferase, a well characterized Rho1 inhibitor^13, 25, 26^, in pre-gastrulating embryos just before the end of cellularization. This timing allowed the C3-transferase to penetrate the cells (because of its low cell-permeability) while not drastically perturbating the early steps of gastrulation (e.g. mesoderm/endoderm invagination). In the second case, we generated maternal/zygotic null mutant embryos for *RhoGEF2* with germline clones (see Methods). *RhoGEF2* encodes a Rho guanine nucleotide exchange factors (RhoGEFs) which is believed to be the main if not the sole GEF activating Rho1 in the medio-apical cortex of embryonic *Drosophila* epithelial cells^16, 27, 28^. In both cases, we achieved a complete loss of medial MyoII pulsatility, resulting in cell and tissue abnormalities (see apical rounding and AJs lowering in Fig. 2b and Supplementary Movie 6). Despite very strong inhibition of MyoII we also noticed that the persistent network was preserved in both C3-transferase injected and *RhoGEF2*^-/-^ embryos (Fig. 2b). Indeed, by measuring the mean levels, we found that, the average medial F-Actin density was only slightly reduced under these inhibitory conditions (Fig. 2c). This shows that the persistent network assembly does not rely on the mechanisms promoting the cortical pulsed contractility.

**Fig. 2.**
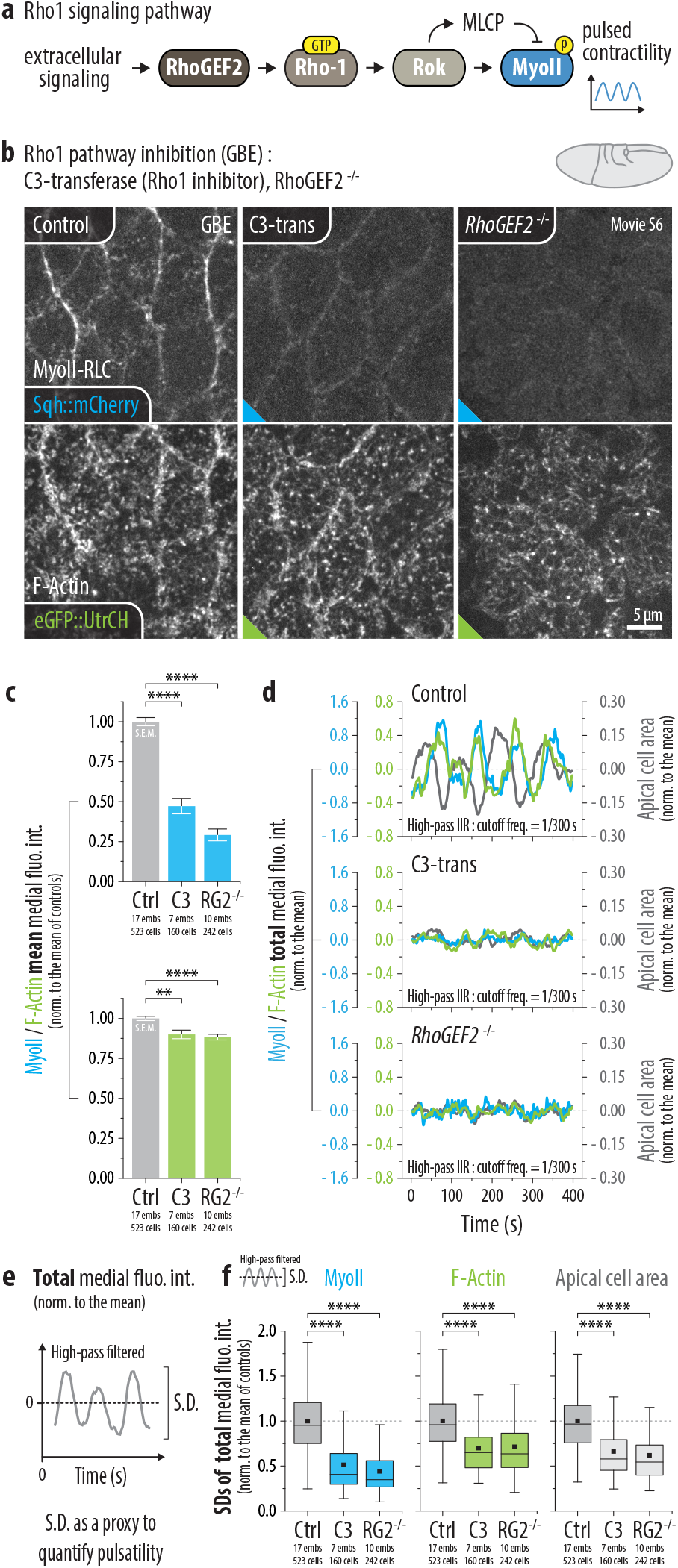
Rho1 pathway inhibition reveals two differentially regulated medio-apical F-Actin sub-populations (GBE). **(a)** Diagram representing the molecular components of the Rho1 signaling pathway. **(b)** Live MyoII-RLC (Sqh::mCherry) and F-Actin (eGFP::UtrCH) localization in ectodermal cells during GBE, in control, C3-transferase injected or *RhoGEF2*^-/-^ mutant embryos. Images represent a single time-point extracted from the Supplementary Movie 6, max-proj. (4 x 0.33 µm). **(c)** Bar plots : mean ± S.E.M. medial MyoII (top panel) and F-Actin (bottom panel) fluo. int. averaged per cell and over time (150 x 3 s). Results are normalized to the mean of controls : water injected (C3-transferase) or WT (*RhoGEF2*^-/-^). Controls are grouped together for a concise display. **(d)** Temporal variations of apical cell area (grey), total medial MyoII-RLC (blue) and F-Actin (green) fluo. int. in a selected cell. Data are high-pass filtered (cutoff freq. 1/300 s) and normalized to the mean. **(e)** Measuring S.D. of high-pass filtered apical cell area or total medial fluo. int. fluctuations as a proxy to quantify pulsatility. **(f)** Box plots : cell averaged S.D. of high-pass filtered total medial MyoII-RLC (left) / F-Actin (middle) fluo. int. and apical cell area (right). Results are normalized to the mean of controls. Box plots (f) : extend from 1^st^ (Q1) to 3^rd^ (Q3) quartile (Q3-Q1 = IQR), whiskers : Q1 or Q3 ± 1.5 x IQR, horizontal lines : medians, black squares : means. Statistical significance (c,f) : two-sample t-test, NS : p > 5E-2, * : p < 5E-2, ** : p < 5E-3, *** : p < 5E-4, **** : p< 5E-5.

We next tested the impact of Rho1 pathway inhibition on F-Actin polymerization during MyoII pulses. To this end, we monitored single cell total fluorescence intensity profiles and quantified the standard deviation as a proxy to measure pulsatility (Fig. 2d,e). Note that we processed these profiles with a high-pass filter to eliminate the low frequency components and isolate the effect of pulses in our measurements (see Methods and Supplementary Fig. 1d). Following this method, we found that, like the apical cell area and MyoII intensity, the fluctuations of medial F-Actin levels were significantly reduced upon Rho1 pathway inhibitions (Fig. 2f). These results demonstrate that, contrary to the persistent network, the pulsatile pool of medial F-Actin is chiefly regulated by Rho1 signaling.

In a second step, we pursued our investigations by focusing on amnioserosa cells during DC and asked whether a differential regulation of the medio-apical F-Actin is also present in this tissue. However, since the C3-transferase is not membrane permeable and *RhoGEF2*^-/-^ embryos were unable to reach such a late embryonic stage due to earlier requirements, we used alternate strategies. We first took advantage of the salt and pepper expression pattern of *engrailed*-GAL4 (en-GAL4) in the amnioserosa to drive the over-expression of a Rho1 dominant negative form, UAS-Rho1N19, in randomly located amnioserosa cells. To identify these over-expressing cells, we recombined the en-GAL4 driver with a fluorescent nuclear marker, UAS-NLS::RFP, to act as a reporter of expression (see Methods and Fig. 3b). We observed that, as expected when MyoII is inhibited, the cells over-expressing Rho1N19 did not undergo pulsed contractility (see yellow ROI in Fig. 3a and Supplementary Movie 7). Strikingly, as in ectodermal cells during germband extension, our quantifications revealed that only the pulsatile pool of F-Actin, and not the persistent network, was perturbed upon Rho1 inhibition (see similar medial F-Actin density in Fig. 3c and reduced medial F-Actin pulsatility in Fig. 3d). These results establish that, in both ectodermal and amnioserosa cells, the medio-apical F-Actin network consists of two independently regulated but entangled pools of filaments. On one hand, a pulsatile pool of F-Actin polymerizing under the control of Rho1 signaling and, on the other hand, a persistent network whose assembly does not depend on this pathway (see diagram Fig. 3h).

**Fig. 3.**
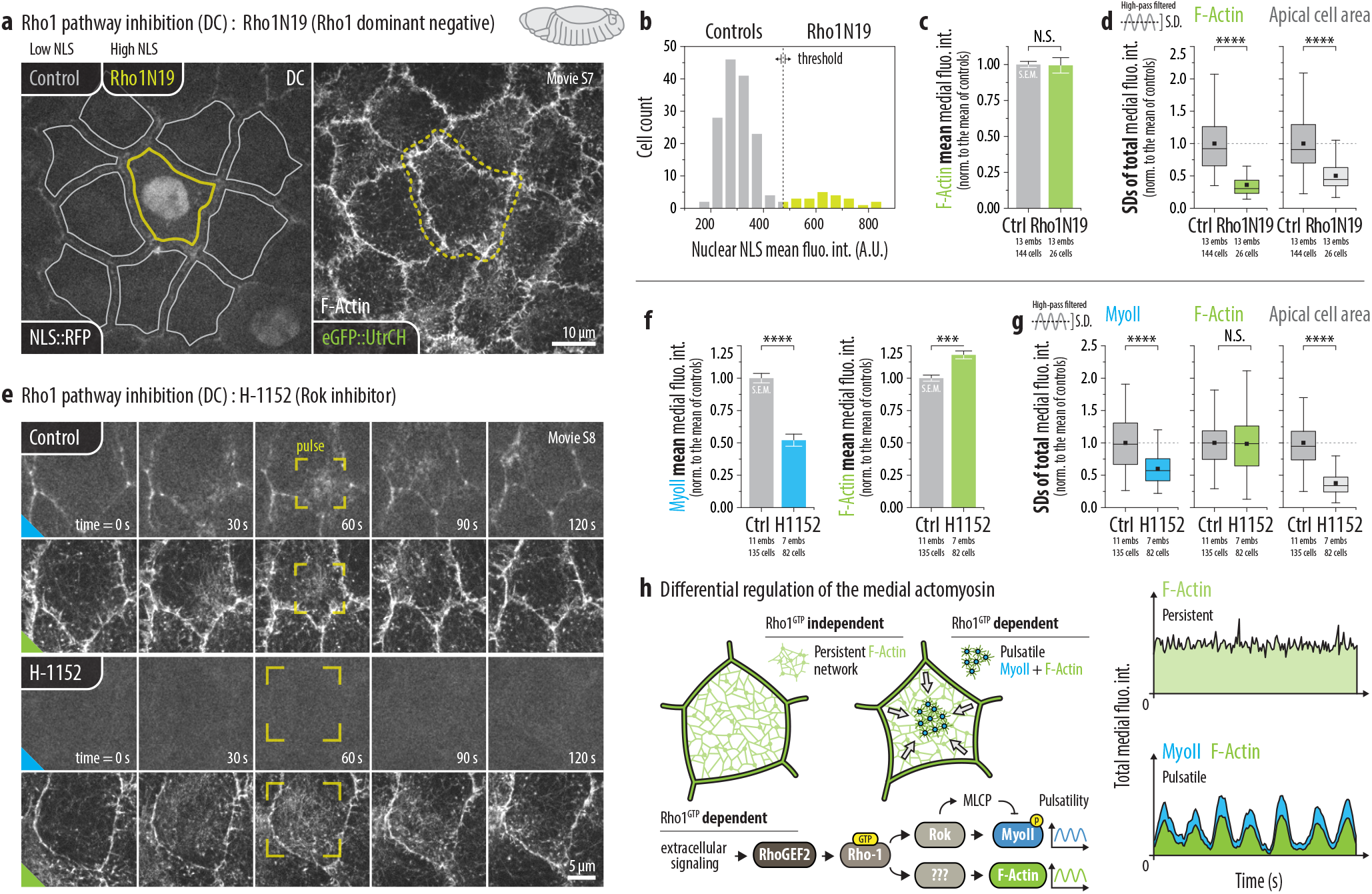
Rho1 pathway inhibition reveals two differentially regulated medio-apical F-Actin sub-populations (DC). **(a)** Live F-Actin (eGFP::UtrCH) localization in amnioserosa cells during DC, in WT (grey outline) or Rho1N19 (yellow outline) overexpressing cells. Images represent a single time-point extracted from the Supplementary Movie 7, max-proj. (4 x 0.33 µm). **(b)** Distribution of measured nuclear NLS::RFP fluo. int. and selected threshold to define the WT (<625) and Rho1N19 overexpressing cells (>625). **(c)** Bar plots : mean ± S.E.M. medial F-Actin fluo. int. averaged per cell and over time (90 or 120 x 10 sec). Results are normalized to the mean of WT cells. **(d)** Box plots : cell averaged S.D. of high-pass filtered (cutoff freq. 1/600 s) total medial F-Actin (left) fluo. int. and apical cell area (right). Results are normalized to the mean of WT cells. **(e)** Live MyoII-RLC (Sqh::mKate2) and F-Actin (eGFP::UtrCH) localization in amnioserosa cells during DC, in control (water injected) and H-1152 injected embryos. Time series : images are extracted from the Supplementary Movie 8, max-proj. (4 x 0.33 µm). The yellow frames show the typical spread of a pulse in these two conditions. **(f)** Bar plots : mean ± S.E.M. medial F-Actin fluo. int. averaged per cell and over time (90 or 120 x 10 s). Results are normalized to the mean of controls. **(g)** Box plots : cell averaged S.D. of high-pass filtered (cutoff freq. 1/600 s) total medial MyoII-RLC (left) / F-Actin (middle) fluo. int. and apical cell area (right). Results are normalized to the mean of controls. **(h)** Diagrams representing the differential regulation of medio-apical actomyosin. A Rho1 independent pathway promotes the persistent F-Actin network assembly (F-Actin baseline) while Rho1 activity underlies both the MyoII and F-Actin pulsatility. Box plots (d,g) : extend from 1^st^ (Q1) to 3^rd^ (Q3) quartile (Q3-Q1 = IQR), whiskers : Q1 or Q3 ± 1.5 x IQR, horizontal lines : medians, black squares : means. Statistical significance (c,d,f,g) : two-sample t-test, NS : p > 5E-2, * : p < 5E-2, ** : p < 5E-3, *** : p < 5E-4, **** : p< 5E-5.

In a last experiment, we wanted to know if the emergence of medial F-Actin pulsatility was dependent on MyoII activation. To do so, we inhibited the Rho associated kinase (Rok) in amnioserosa cells, by injecting the cell-permeable H-1152 compound^13^ in embryos at the DC stage. Interestingly, while these injections were successful to inhibit MyoII pulses and the apical cell contractility, our quantifications revealed that both the persistent network and the pulsatile polymerizations of F-Actin were preserved in this condition (Fig. 3e-g and Supplementary Movie 8). The dynamics of F-Actin pulses was however modified upon Rok inhibition, with pulses tending to last longer and be larger than in controls (see yellow frames in Fig. 3e). Although these changes likely reflect the role of MyoII contractility in shaping pulses of F-actin polymerization, we conclude that the emergence, *per se*, of such pulses does not rely on Rok/MyoII activity itself. Overall Rho1 signaling is responsible for both MyoII and F-Actin pulsatility but uses different downstream effectors (see diagram Fig. 3h).

### The Frl formin promotes the persistent F-Actin network assembly

While our results clearly demonstrated the role of Rho1 signaling in the emergence of medial F-Actin pulsatility, the mechanisms underlying assembly of the medial apical actin network in the early germband and in the amnioserosa are unknown. We performed an shRNA screen to identify the factor(s) responsible for the persistent network assembly. Since the network density is relatively low, we focused our efforts on the actin nucleators of the formin family, known to promote the assembly of sparse F-Actin networks^17^. By first looking at the actomyosin dynamics in amnioserosa cells, we noticed that a down-regulation of the Frl/Fmnl formin leads to a clear loss of medial F-Actin density in the time interval between two MyoII/F-Actin pulses (see “no pulse” vs “pulse” in Fig. 4a and Supplementary Movie 9). This result led us to consider Frl as a good candidate for the persistent network assembly and to further investigate its function in both ectodermal (GBE) and amnioserosa cells (DC).

**Fig. 4.**
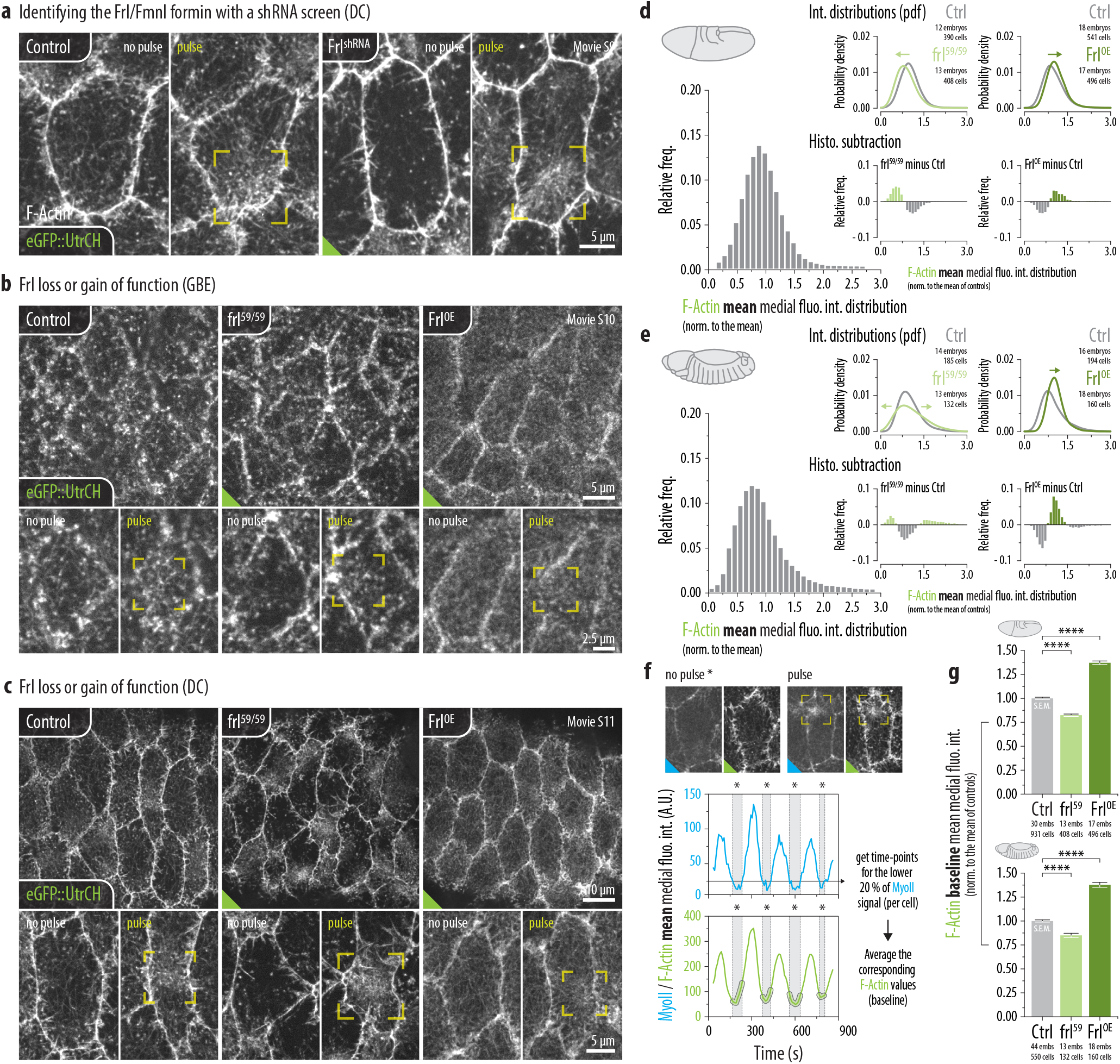
The Frl/Fmnl formin promotes the persistent F-Actin network assembly. **(a,b,c)** Live F-Actin (eGFP::UtrCH) localization in ectodermal (GBE) or amnioserosa cells (DC) in the following conditions : WT, Frl^shRNA^ (shRNA against Frl), *frl*^59/59^ (frl null mutant) or Frl^OE^ (Frl overexpression). Images represent either a time-point between two pulses (“no pulse”) or during a pulse (“pulse”) and are extracted from : (a) Supplementary Movie 9 (DC, WT vs Frl^shRNA^); (b) Supplementary Movie 10 (GBE, WT vs *frl*^59/59^ vs Frl^OE^); (c) Supplementary Movie 11 (DC, WT vs *frl*^59/59^ vs Frl^OE^). The yellow frames show the pulse localization. **(d,e)** Mean medial F-Actin fluo. int. distributions in Frl loss or gain of function during GBE (d) or DC (e). Main bar plots : distributions for WT embryos. Probability density functions (pdf) and histograms subtraction : distributions comparison and relative bins enrichment between WT and *frl*^59/59^ (left) or WT and Frl^OE^ (right). All distributions are normalized to the mean of controls. **(f)** Measuring the persistent network density (F-Actin baseline) by averaging the mean medial F-Actin fluo. int. during the 20% lowest mean medial MyoII fluo. int. time-points. **(g)** Bar plots : mean ± S.E.M. medial F-Actin baseline averaged per cell during GBE (top panel) or DC (bottom panel). Results are normalized to the mean of WT cells. Statistical significance (g) : two-sample t-test, NS : p > 5E-2, * : p < 5E-2, ** : p < 5E-3, *** : p < 5E-4, **** : p< 5E-5.

First, to confirm our loss of function phenotype, we produced a null allele of frl (*frl*^59^) by adopting a CRISPR deletion strategy (see Methods). While this allele revealed to be semi-lethal and semi-sterile, we succeeded to cross homozygote parents (*frl*^59/59^) and obtained maternal/zygotic null embryos. In a complementary approach, we also studied the effect of a Frl gain of function by overexpressing a UAS-Frl^wt^ construct using a 67-GAL4 driver (condition referred as Frl^OE^). In both tissues, we observed that the expression level of Frl affects the persistent actin network density: the panels in Fig. 4b,c indicated “no pulse” show the reduced network density in *frl*^59/59^ mutants and the increased network density in Frl^OE^ embryos (see also Supplementary Movies 10,11). Furthermore, as we will further describe below, we also found that Frl influences the pulsatile pool of F-Actin, especially in amnioserosa cells, by reducing the amplitude of pulses (the more Frl, the weaker pulses). In light of this observation, we carefully designed our quantification methods to measure the persistent network density, considering that both the persistent network and the pulsatile pool account for the medial F-Actin levels and that Frl affects these two sub-populations. We first looked at the distribution of the medial F-Actin density and compared the relative changes between conditions (Fig. 4d,e). Modulating the Frl expression induced a shift in the medial F-Actin density distributions, towards lower values in *frl*^59/59^ mutants and towards higher values in Frl^OE^ embryos (Fig. 4d,e, arrows above the graphs), consistent with a decrease or an increase of the persistent network density.

Next, to address how Frl levels influence the persistent network density independent of its contribution to the pulsatile pool of F-Actin, we monitored single cell mean fluorescence intensity profiles for the MyoII/F-Actin and selected the 20 % time-points for which the MyoII signal was the lowest (Fig. 4f). Considering these time-points as inter-pulses period, we then measured and averaged the corresponding mean F-Actin levels (F-Actin baseline) and observed that, in both tissues, lowering the Frl levels induced a reduction of medial F-Actin density while overexpressing Frl produced the opposite effect (Fig. 4g). These results confirm that Frl plays a pivotal role in the persistent network assembly in ectodermal and amnioserosa cells.

### Frl antagonizes Rho1-induced medial pulsed contractility

To better characterize how modulating the Frl levels also influences the pulsatile pool of F-Actin and the medio-apical contractility in general we monitored the medio-apical actomyosin levels and the apical cell area fluctuations by looking at the standard deviations of high-pass filtered single cell profiles (Fig. 5a-d and Supplementary Movies 12,13). In amnioserosa cells, the loss of the Frl (*frl*^59/59^) resulted in a clear increase of both the MyoII/F-Actin and the apical cell area fluctuations. In contrast, its overexpression (Frl^OE^) produced the opposite effect (Fig. 5d). We observed similar tendencies in ectodermal cells, albeit to a lower extent (Fig. 5c). Taken together these quantifications showed that, beyond its role in the persistent network assembly, Frl counteracts the medial actomyosin pulsatility and the apical cell surface deformation.

**Fig. 5.**
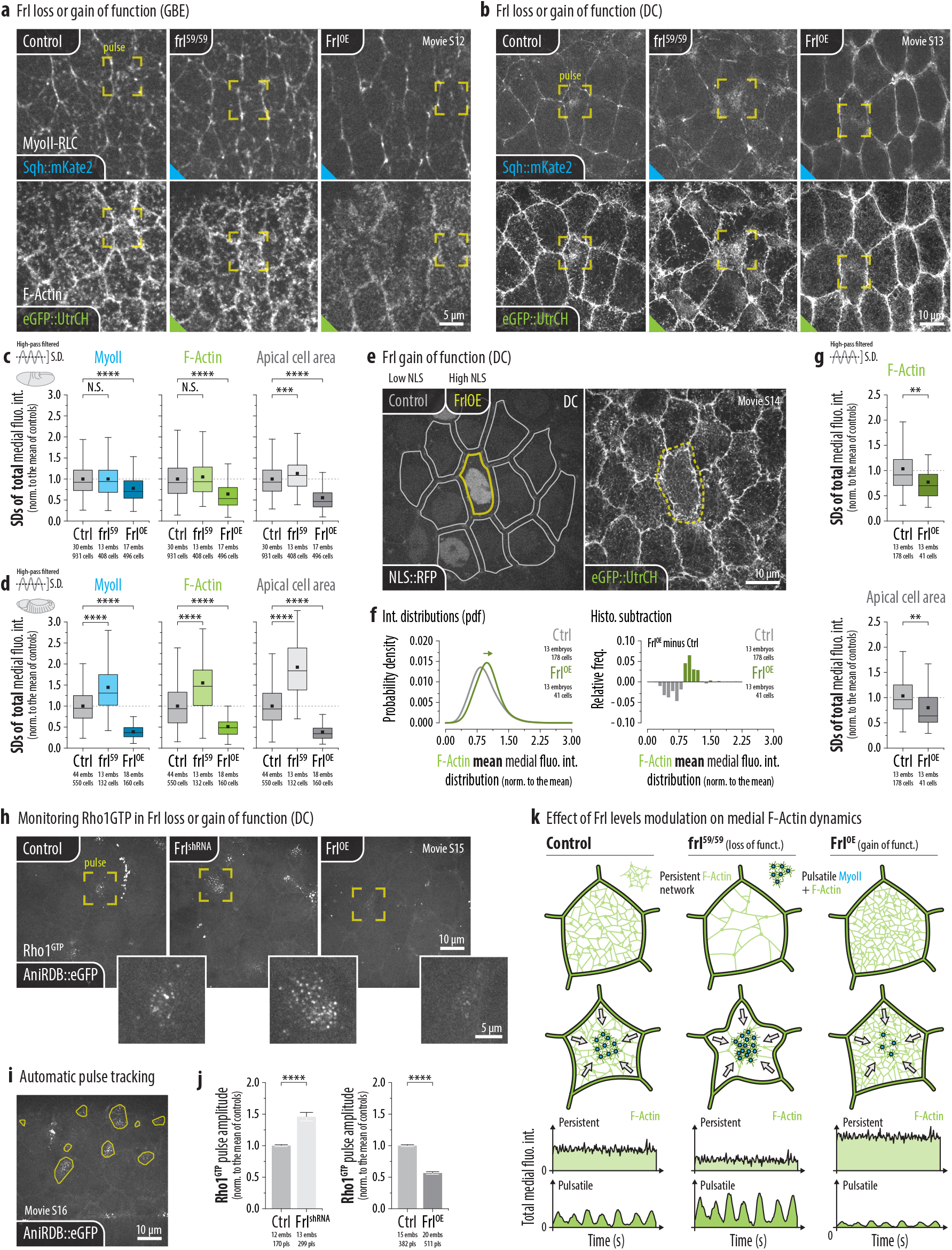
Frl antagonizes the Rho1-induced medial pulsed contractility. **(a,b)** Live MyoII-RLC (Sqh::mKate2) and F-Actin (eGFP::UtrCH) localization in ectodermal (GBE) or amnioserosa cells (DC), in WT, *frl*^59/59^ or Frl^OE^ embryos. Images represent a single time-point extracted from the Supplementary Movie 12 (GBE) or 13 (DC), max-proj. (4 x 0.33 µm). The yellow frames show the pulse localization. **(c,d)** Box plots : cell averaged S.D. of high-pass filtered (cutoff freq. 1/300 s for GBE and 1/600 s for DC) total medial MyoII-RLC (left) / F-Actin (middle) fluo. int. and apical cell area (right). Results are normalized to the mean of controls. **(e)** Live F-Actin (eGFP::UtrCH) localization in amnioserosa cells during DC, in WT (grey outline) or Frl (yellow outline) overexpressing cells. Images represent a single time-point extracted from the Supplementary Movie 14, max-proj. (4 x 0.33 µm). **(f)** Mean medial F-Actin fluo. int. distributions in WT or Frl overexpressing amnioserosa cells during DC. Probability density functions (pdf) and histograms subtraction : distributions comparison and relative bins enrichment between WT and Frl^OE^. All distributions are normalized to the mean of controls. **(g)** Box plots : cell averaged S.D. of high-pass filtered (cutoff freq. 1/600 s) total medial F-Actin (top) fluo. int. and apical cell area (bottom). Results are normalized to the mean of controls. **(h)** Live Rho1GTP (AniRBD::eGFP) localization in amnioserosa cells during DC, in WT, Frl^shRNA^ or Frl^OE^ embryos. Main images : single time-point extracted from the Supplementary Movie 15, max-proj. (4 x 0.33 µm). Inserted images : selected zoomed on a pulse (see yellow frames, main images). **(i)** Automatic segmentation and tracking of Rho1GTP pulses (see Methods and Supplementary Movie 16). **(j)** Bar plots : mean ± S.E.M. Rho1GTP pulse amplitude calculated on the total int. of segmented ROIs. Results are normalized to the mean of WT embryos. **(k)** Diagrams representing how modulating the Frl levels influence the pulsed contractility. In Frl loss of function (*frl*^59/59^, Frl^shRNA^) the persistent network density is reduced while the MyoII/F-Actin pulsatility and apical cell contractility are increased. Overexpressing Frl (Frl^OE^) induces opposite effects. Box plots (c,d,g) : extend from 1^st^ (Q1) to 3^rd^ (Q3) quartile (Q3-Q1 = IQR), whiskers : Q1 or Q3 ± 1.5 x IQR, horizontal lines : medians, black squares : means. Statistical significance (c,d,g,j) : two-sample t-test, NS : p > 5E-2, * : p < 5E-2, ** : p < 5E-3, *** : p < 5E-4, **** : p< 5E-5.

We further tested whether the effect on pulsation is cell autonomous of whether it involves cell interactions. As we did with Rho1N19 (see Method and Fig. 3a), we overexpressed Frl using the *engrailed*-GAL4 driver to see if the Frl effect on the pulsed contractility can be obtained in isolated amnioserosa cells (Fig. 5e and Supplementary Movie 14). We observed that cells overexpressing Frl displayed an increased persistent network density (see distribution shift in Fig 5f) and a reduced F-Actin pulsatility and apical cell area fluctuations (Fig. 5g). This experiment confirmed that Frl antagonizes the medial actomyosin pulsatility and that this effect occurs cell autonomously.

We then asked whether Frl influences the actomyosin pulsatility by modulating the level of Rho1 activity (Rho1GTP) during pulses. To address this question, we monitored the medio-apical localization of the Rho1 binding domain of anillin fused to eGFP (AniRBD::eGFP) in amnioserosa cells. This construct acts as a sensor to follow Rho1 activity *in vivo* since it specifically binds to the GTP bound form of Rho1^13, 29^. We next combined the sensor with a 67-Gal4 driver to either overexpress a UAS driven shRNA (Frl^shRNA^) or the UAS-Frl^wt^ construct (Frl^OE^) in embryos (Fig. 5h and Supplementary Movies 15). We observed and measured (see Methods, Supplementary Fig. 2a and Supplementary Movies 16) that reducing the Frl levels enhanced the amplitude of AniRBD::eGFP pulses while overexpressing the formin led to the opposite effect (Fig. 5i,j). This allowed us to conclude that Frl significantly affects the apical cell contractility by modulating the levels of Rho1 activation.

Overall, we conclude that tuning the Frl levels act as a switch between two distinct modes of medio-apical contractility. At zero/low level of Frl (*frl*^59/59^ and Frl^shRNA^) the persistent network is seriously weakened and the actomyosin pulses have an increased amplitude. Consequently, cells undergo more pronounced cell shape changes. In sharp contrast, the cells overexpressing Frl are more static, showing a low contractility and a dense persistent network (see diagram Fig. 5k).

### Cellular and tissue scale effects of Frl loss or gain of function

We next wanted to know if changes in cellular behavior observed following the modulation of the Frl levels can in turn influence the overall tissue dynamics. To answer this question, we first monitored ectodermal cells undergoing convergent extension (GBE) by segmenting cells individually (Fig. 6a and Supplementary Movie 17). Since convergent extension is driven by cell intercalation (T1 events) and that cell intercalation itself is powered by MyoII pulses^7, 30, 31^, we tracked T1 events and monitored cell area fluctuations as a read out of pulsatility. Our measurements revealed that ectodermal cells intercalate and fluctuate more in *frl*^59/59^ mutants and less in Frl^OE^ than in control embryos (see inserted time projection in Fig. 6a and quantifications in Fig. 6b,c). In general, the occurrence of T1 events and the intensity of apical cell area fluctuations are positively correlated across conditions (Pearson coefficient = 0.61, sig. 6.1 x 10^-6^, Fig. 6d). This suggests that active fluctuations favor T1 events and that Frl, by tuning these fluctuations locally, can impacts the germband dynamics at the tissue scale. Interestingly, while increasing cell intercalation (*frl*^59/59^) does not speed up germband extension reducing the occurrence of T1 events (Frl^OE^) slows it down (assessed by following the posterior mid gut in DIC movies, see Fig. 6e,f and Supplementary Movie 18). This observation is consistent with previous reports that reducing cell intercalation, using other mutant conditions (e.g. eve mutants, Toll2, 6, 8 RNAi), impaired germband extension^32, 33^.

**Fig. 6.**
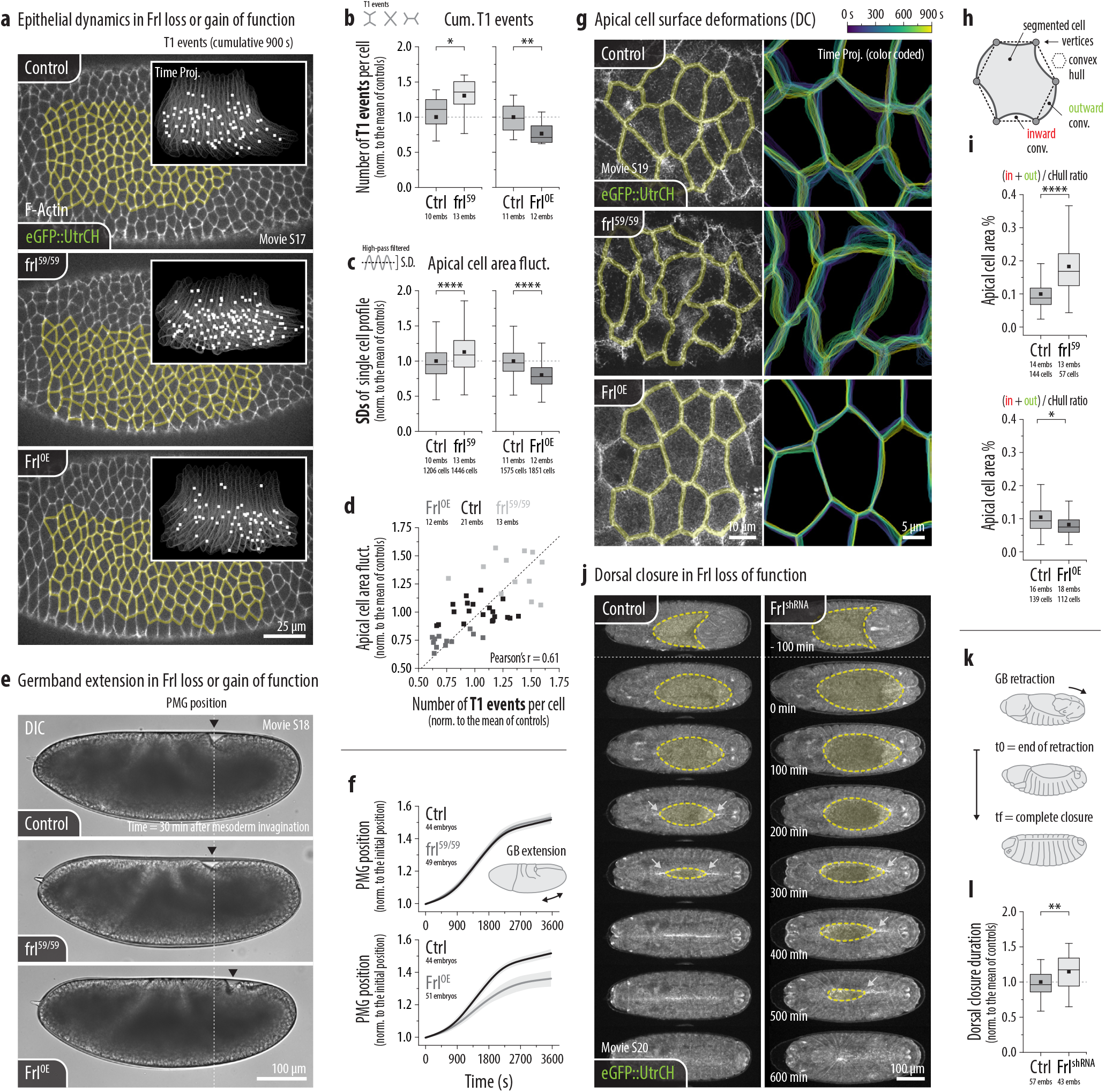
Cellular and tissue scale effects of Frl loss or gain of function. **(a)** Live F-Actin (eGFP::UtrCH) in ectodermal cells revealing epithelial dynamics during GBE in WT, frl^59/59^ or Frl^OE^ embryos. Main images : single time-point extracted from the Supplementary Movie 17. The yellow cell outlines show the result of the automatic cell segmentation procedure. Inserted images : time-projection (900 s) of the cell outlines (grey) and the recorded T1 events (white squares). **(b)** Box plots : number of T1 events per cell for the duration of the movies (900 s). Results are normalized to the mean of WT embryos. **(c)** Box plots : cell averaged S.D. of high-pass filtered (cutoff freq. 1/600 s) apical cell area. Results are normalized to the mean of WT. **(d)** Scatter plots : apical cell area fluctuations as a function of the number of T1 events per cell in WT, frl^59/59^ or Frl^OE^ condition. The dashed line shows a linear fit with a Pearson’s correlation coefficient r = 0.61. **(e)** Live DIC movies of embryos undergoing GBE in WT, frl^59/59^ or Frl^OE^ condition. Images represent a single time-point 30 min after the onset of gastrulation and are extracted from the Supplementary Movie 18. The black arrowheads show the posterior mid-gut (PMG) position. **(f)** Line plots : PMG position ± S.D. over time revealing the progression of the germband extension. Data are normalized to the initial PMG position. **(g)** Live F-Actin (eGFP::UtrCH) in amnioserosa cells during DC in WT, frl^59/59^ or Frl^OE^ embryos. Left images : single time-point extracted from the Supplementary Movie 19, max-proj. (4 x 0.33 µm). The yellow cell outlines show the result of the automatic cell segmentation procedure. Right images : color coded time-projection (900 s) of the cell outlines. **(h)** Measuring cell shape irregularity by comparing the convex hull (connecting vertices) and the segmented apical cell surface. **(i)** Box plots : ratio between the surface of the inward + outward regions and the surface occupied by the convex hull. **(j)** Live F-Actin (eGFP::UtrCH) low magnification imaging of embryos undergoing DC in WT or Frl^shRNA^ condition. Time series : images are extracted from the Supplementary Movie 20 at the indicated time-points. The yellow dashed lines and the surrounded regions shows the amnioserosa cells at the surface of the embryo. **(k)** Schematic representation of the DC process. **(l)** Box plots : DC duration normalized to the mean of WT embryos. Box plots (b,c,i,l) : extend from 1^st^ (Q1) to 3^rd^ (Q3) quartile (Q3-Q1 = IQR), whiskers : Q1 or Q3 ± 1.5 x IQR, horizontal lines : medians, black squares : means. Statistical significance (b,c,i,l) : two-sample t-test, NS : p > 5E-2, * : p < 5E-2, ** : p < 5E-3, *** : p < 5E-4, **** : p< 5E-5.

In a second step, we studied how modulating the Frl levels affects tissue dorsal closure. As we have seen before, the Frl mutant phenotype is particularly pronounced in amnioserosa cells, as a lack of Frl induces a strong increase of actomyosin pulsation and apical cell area fluctuations (see Fig. 5). We also noticed that modulating the Frl levels changes the way cells deform, with cells in the *frl*^59/59^ condition being more irregularly shaped than in control and, even more so, than in Frl overexpressing embryos (see time projected cell boundaries in Fig. 6g and Supplementary Movie 19). Using convex hull (see Methods and Fig. 6h) we measured that cells present indeed more inward and outward convolutions in the Frl loss of function than in the other conditions (see Fig. 6i). Since amnioserosa cells provide the main forces necessary to complete dorsal closure^34, 35^, we assumed that these local modification of the cellular behavior could underlie dorsal closure defects. We therefore imaged embryos at a lower magnification and measured the time elapsed between the end of germband retraction and the complete lateral epidermis closure (Fig. 6j,k and Supplementary Movie 20). By comparing control and Frl shRNA expressing embryos (Frl^shRNA^), we found that down-regulating Frl induces tissue scale defects and slows down closure by ∼15% (Fig. 6l). It appeared that in many Frl^shRNA^ embryos dorsal closure occurs only from the posterior side and not from both posterior and anterior sides like in controls (see white arrows in Fig. 6l). We suggest that despite reinforcing pulsatility, the reduction of Frl levels (Frl^shRNA^) increases cell deformability and could also impair the capability of amnioserosa cells to efficiently propagate contractile forces across junctions and pull on the lateral ectoderm. In other words, Frl may be required for generate an effective large-scale tissue tension. This could be explained by the fact that reducing the persistent network density leads to an overall decrease of apical cortex stiffness, known to increase drastically right at the onset of dorsal closure in WT embryos^36^.

### The persistent network promotes the propagation of MyoII-induced contractile forces

This led us to address how the persistent network the propagation of contractile forces in the tissue. To that end, we focused on the amnioserosa cells, where the modulation of Frl levels had the strongest phenotypic consequences. We observed that pulses tend to contract the whole apical surface in control cells while, in *frl*^59/59^ cells, they exert contractile forces mostly on their close surroundings where pulsatile F-Actin is dense (Fig. 7a). This is especially striking when a pulse travels through the mutant cells, only contracting nearby AJs (Supplementary Movie 21). As a result, pulses in the *frl*^59/59^ mutant generally do not affect the distant parts of the cell, except when some sparse radial filaments, emitted by the pulse itself, connect to the AJs (Fig. 7b). This can lead to surprising deformation dynamics, with distant AJs expanding while nearby AJs are contracting (Fig. 7a). Together, these results suggest that F-Actin supports the transmission of contractile forces between pulses and AJs. In control cells, the homogeneous persistent network distributes evenly contractile forces to the periphery while, in *frl*^59/59^ cells, the lack of this network leads to a heterogenous transmission of these forces (Fig. 7c). This is consistent with our previous quantification of increased deformation heterogeneity in *frl*^59/59^ cells (Fig. 6j-l).

**Fig. 7.**
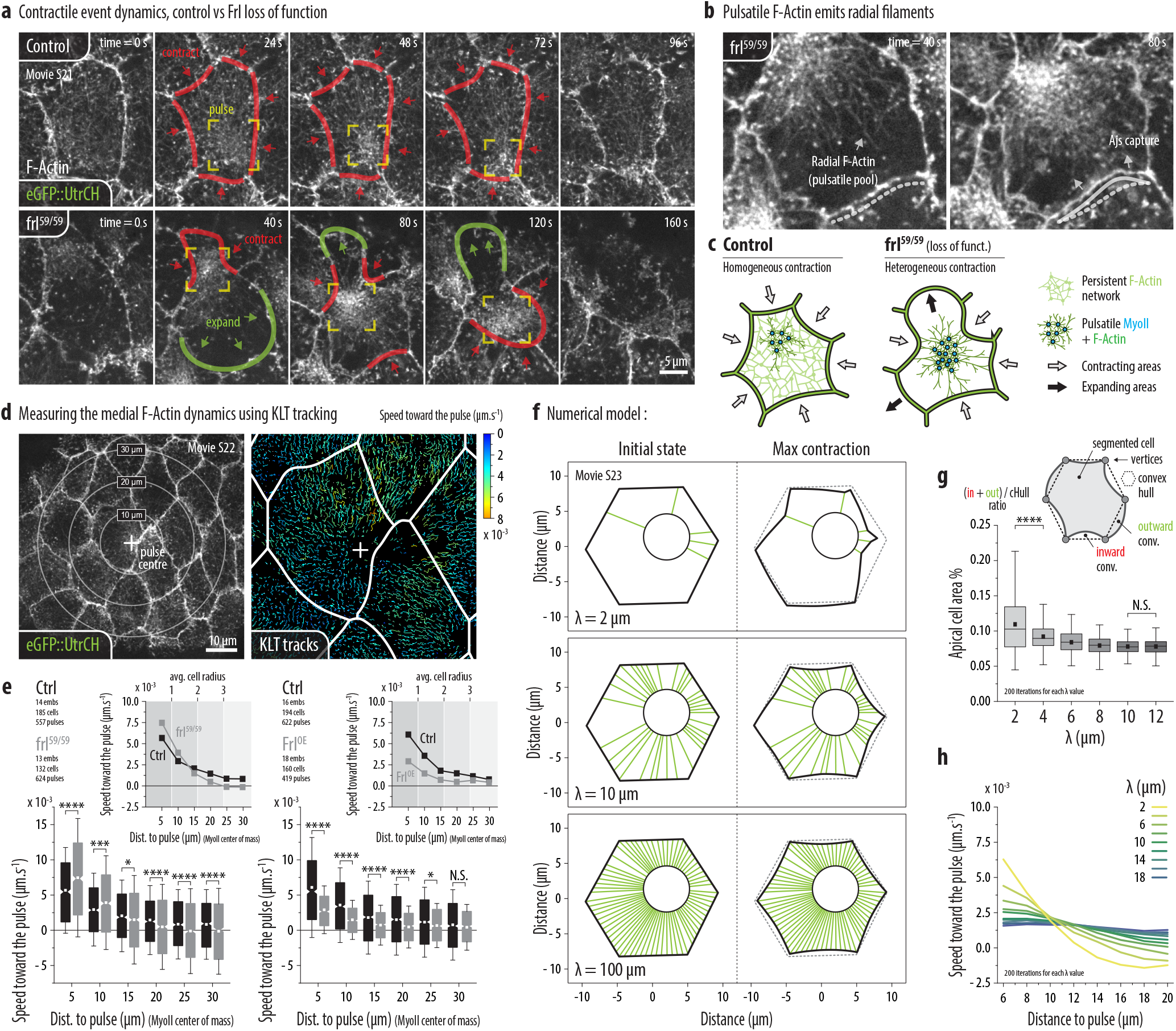
The persistent network promotes the propagation of MyoII-induced contractile forces. **(a)** Live F-Actin (eGFP::UtrCH) localization in amnioserosa cells during DC comparing the dynamic of a pulsatile event in a WT and a *frl*^59/59^ cell. Time series : images are extracted from the Supplementary Movie 21 at the indicated time-points, max-proj. (4 x 0.33 µm). The yellow frames show the pulse localization, the red and green outlines/arrows show, respectively, the contracting and the expanding parts of the cell. **(b)** Zoomed view on a contracting *frl*^59/59^ cell displaying radial F-Actin filaments emanating from the pulse core and capturing the AJs. **(c)** Schematic representation of a contracting cell in WT or *frl*^59/59^ condition. **(d)** Measuring the propagation of pulsatile contractility by following discrete apical F-Actin structures using a KLT tracking procedure. Left image : concentric circles showing the distance from a pulse (yellow cross). Right image : single time-point extracted from the Supplementary Movie 22, displaying the color-coded speed of KLT tracked structures toward the pulse centre. The white lines show the segmented cell boundaries. **(e)** Box plots : averaged speed toward the pulse centre of KLT tracked structures as a function to the distance to the pulse in WT vs *frl*^59/59^ (left) or WT vs Frl^OE^ (right) amnioserosa cells. Data are binned as indicated (distance ± 2.5 µm, e.g. the 5 µm bin contains all tracks within a 2.5 to 7.5 µm distance to the pulse). Line plots : speed toward the pulse as a function to the distance to the pulse averaged per bin. Each shade of grey in the background represents the typical size of an amnioserosa cell radius (~ 8 µm). **(f)** Representative simulations for different values of *λ* (Supplementary Movie 23). Left panels depict the initial condition, right panels depict the cell state upon maximal contraction. Green segments indicate that a boundary element is connected to the pulse. The pulse position is the same in the three examples. **(g)** Diagram : Measuring cell shape irregularity by comparing the convex hull (connecting vertices) and the segmented apical cell surface. Box plots : ratio between the surface of the inward + outward regions and the surface occupied by the convex hull (see Fig. 6k-l) upon maximal deformation. 200 iterations were performed for each value of *λ*. For each iteration, the pulse position is chosen randomly within the cell. **(h)** Line plots : Averaged speed towards the pulse vs. distance to the pulse during the contraction phase. Averages were performed from 200 iterations for each value of *λ*. For each iteration, the pulse position is chosen randomly within the cell. Box plots (e,g) : extend from 1^st^ (Q1) to 3^rd^ (Q3) quartile (Q3-Q1 = IQR), whiskers : S.D. (e) or : Q1 or Q3 ± 1.5 x IQR, horizontal lines : medians, black squares : means. Statistical significance (e,g) : two-sample t-test, NS : p > 5E-2, * : p < 5E-2, ** : p < 5E-3, *** : p < 5E-4, **** : p< 5E-5.

Next, to further characterize how the heterogeneity of F-Actin distribution impacts pulsed contractility, we measured the distance at which contractile forces propagate within the cell and tissue. To do so, we performed a KLT analysis measuring the speed at which tracked apical F-Actin structures move towards the pulse as a function of their distance to the pulse. (see Fig. 7d, Methods, Supplementary Fig. 2b,c, and Supplementary Movie 22). After binning results according to distance, we were able to produce speed propagation curves and compare measurements between conditions (Fig. 7e). In all cases, speeds were higher near the pulse and decayed gradually with the distance However, we observed that the decay length is strikingly shorter in *frl*^59/59^ cells that in controls, despite the fact that we actually measured higher speeds at close range in mutant cells. This results in a crossover between the *frl*^59/59^ and control curves. Note however that such a crossover was not present between Frl^OE^ and control cells, although contraction amplitudes were different. These results indicate that the homogeneous persistent network promotes propagation of contractile forces at longer range.

We first reasoned that the F-Actin network behaves as a continuous material and the shorter decay length observed in *frl*^59/59^ cells might result from a reduction of the so-called hydrodynamic length. Indeed, mechanical information typically does not propagate beyond this length, which is the distance within reach before internal dissipation occurs^23^. In a viscoelastic scenario, this distance increases with stiffness. This is consistent with our observations, as frl^59/59^ cells display an overall reduction of F-Actin density, which is likely to reduce the effective stiffness. However, the time required to propagate over the hydrodynamic length is typically the dissipation timescale. In F-Actin networks, dissipation is highly influenced by filaments turnover which have been measured to occur over timescales no shorter than 10s, and possibly more^37–39^. We did not observe such contraction delay between local and distant regions of the cell (Supplementary Fig. 2d). Within the propagation range, contractions occur almost simultaneously. This rule out the hypothesis wherein Frl tunes the propagation range by affecting the hydrodynamic length.

We next reasoned that heterogeneity of connectivity to distant AJs might reduce the propagation range of contractile forces. To test this hypothesis, we designed a numerical model in which the cell boundary is discretized (see Methods and Supplementary Fig. 2e). The pulse position is chosen randomly, and a contraction force directed towards the pulse is applied to elements of the cell boundary. To consider the variable connectivity of the network, we assume that the probability for an element of the boundary to be connected to the pulse (and thus to directly undergo the contractile force) decays on a length scale *λ* with the distance *d* to the pulse, *p*∼*e^−d/λ^*. This view is reminiscent of percolation systems in which the size of connected subregions increases with density^40^. Clearly, *λ* should be small in a low connectivity situation (*frl*^59/59^ cells), when there is no persistent network. Yet even in this scenario, a distant element has a non-zero probability to be connected to the pulse, consistent with our observations of sparse radial filaments in *frl*^59/59^ cells. In contrast, *λ* should increase when the network density increases (WT and Frl^OE^ cells), so that when *λ* is large enough, the whole cell boundary is eventually connected to the pulse.

First, our model recapitulates the cell contraction phenotypes observed in the different conditions (Fig. 7f and Supplementary Movie 23). When *λ* is small, force is mostly transmitted to proximal boundary regions, and connection to distant AJs is very sparse. As a result, contraction is heterogeneous, and possibly leads to local expansion in distant regions due to area constraints. When *λ* increases, both connectivity and contraction become more homogeneous. Consequently, *λ* directly impacts the shape of contracting cells, and simulations with a small value of *λ* lead to more convoluted cell shapes, as previously observed for *frl*^59/59^ cells (compare simulations in Fig. 7g and actual *in vivo* measurements in Fig. 6j-l). In simulations, we also observe that variability is much higher when *λ* is small (Fig. 7g). This directly results from the stochastic nature of the low connectivity regime, in which only a random subset of boundary elements is connected. The more boundary elements are connected to the pulse, the more reproducible the deformation pattern becomes. Interestingly, variability was also much higher in *frl*^59/59^ cells than in control cells (Fig. 6i), which further indicates sparse, random connectivity to AJs. Second, we averaged the movement towards the pulse as a function of the distance to the pulse for a wide range of *λ* (Fig. 7h). This recapitulated qualitatively the observations of Fig. 7e, in particular the crossover between propagation curves at low *λ*. Interestingly, when *λ* increases beyond cell size, connectivity eventually saturates, and no more crossovers occur between curves of high *λ*. This is consistent with our results when comparing the WT and Frl^OE^ cells (Fig. 7e).

Overall, our results indicate that the persistent F-Actin network acts as a support for the robust transmission of contractile forces. Absence of this network yields sparse connectivity, which affects homogeneous force transmission to the cell boundaries, and reduces the propagation range of contractile forces.

## Discussion

While most studies concerning the pulsed cortical contractility focused on deciphering the mechanisms underlying the emergence of MyoII pulsatility, we focused here on how the cortical F-Actin influences this process in embryonic *Drosophila* epithelial cells. We showed that, in both ectodermal (GBE) and amnioserosa cells (DC), the medio-apical cortex consists of two differentially regulated, but entangled subpopulations of actin filaments. These two populations share the same sub-cellular localization but undergo distinct spatio-temporal dynamics and influence the pulsed cortical contractility in a different way. The pulsatile F-Actin, together with MyoII, promotes local cell deformations while the persistent homogeneous network ensures homogeneous connectivity between pulses and the AJs and hence spatial propagation of deformation. We identified the Frl/Fmnl formin as a critical nucleator promoting the persistent network assembly. This constitutes a new role for the Frl/Fmnl formin since so far it has been mainly described as participating to the lamellipodia/filopodia formation^41–44^. It would be interesting to know if, like in other systems, Frl is regulated by either Cdc42^41, 44, 45^ or Rac1^46^ to promote the persistent network assembly. Furthermore, it is likely that other formins participate to the persistent network assembly, especially in the germband since the lack of Frl (frl^59/59^) only partially reduces the network density (see Fig. 4b). The DAAM formin would constitute a first good candidate since it has been shown that DAAM and Frl cooperate during axon growth in the mushroom bodies of *Drosophila*^45^.

Although the pulsatile and persistent actin networks show different dependencies on Rho1 activity, we reported that Frl antagonizes the medio-apical Rho1 dependent contractility. Further work is required to identify the crosstalk mechanisms operating between Frl and the pulsed contractility. To this end, it will be interesting to draw from previous studies reporting that the F-Actin can negatively feedback on Rho1 activation^11, 15^. Indeed, it is possible that, like in the *C.elegans* zygote, some Rho1 inhibitors (e.g. RhoGAPs) bind to cortical F-Actin in our systems. Consequently, modifying the persistent network density, through Frl loss or gain of function, could in turn modulate the levels of apical Rho1 activation. It has also been shown that advection acts as a positive feedback for pulsatility, by increasing the local concentration of upstream regulators (e.g. Rho1 and Rok)^13^. It will therefore be interesting to study how the persistent network influence advection and how lowering/increasing the network density affects this feedback mechanism.

Our data also revealed that modulating Frl levels has an impact on epithelial dynamics at the cellular and tissue scale (see Fig. 6). Although this is probably due in part to the effect of Frl on the medial actomyosin pulsatility, we designed a series of analysis to understand how the persistent network may influence the pulsed contractility in mechanical terms. It was suggested that the medio-apical F-Actin acts as a scaffold to transmit contractile forces to the AJs and, by extension, to the surrounding tissue^7, 47–49^. Our results revealed that the persistent network does indeed play a key role in this process by promoting the uniform distribution and the propagation of contractile forces at longer range. We also devised a numerical model recapitulating qualitatively our experimental measurements and providing solid evidences arguing that Frl influences epithelial dynamics through the persistent network assembly, independently from its effect on the actomyosin pulsatility.

Overall, this work sheds new light on how the cortical F-Actin layer assembles *in vivo* and how its dynamic organization influences MyoII-induced stress propagation. Our findings echo to previous experimental and theoretical studies demonstrating that the F-Actin network, through its cross-linking state^50–53^, the length of its filaments^22^ or its turnover^48^ can mediate the amplitude and the length scale at which cortical stresses propagate. We showed here that the F-Actin cortex is composed of differentially regulated sub-populations of filaments influencing its mechanical properties, distinctly, namely contraction and spatial propagation of cortex deformation. Tissue morphogenesis requires interaction between different cellular and tissue level deformation whose propagation in space and time are little understood. It will be important to unravel how cells may tune in different stages of development or in different tissues these properties. Our work suggests that actin network regulation is an important part of this regulatory process.

## Supporting information

Movie-S1.avi

Movie-S2.avi

Movie-S3.avi

Movie-S4.avi

Movie-S5.avi

Movie-S6.avi

Movie-S7.avi

Movie-S8.avi

Movie-S9.avi

Movie-S10.avi

Movie-S11.avi

Movie-S12.avi

Movie-S13.avi

Movie-S14.avi

Movie-S15.avi

Movie-S16.avi

Movie-S17.avi

Movie-S18.avi

Movie-S19.avi

Movie-S20.avi

Movie-S21.avi

Movie-S22.avi

Movie-S23.avi

## Acknowledgements

We are grateful to József Mihály (Biological Research Centre, HAS, Szeged, Hungary) and Andreas Jenny (Albert Einstein College of Medicine, The Bronx, NY, USA) for providing fly stocks. We thank members of the Lecuit and Lenne groups for stimulating discussions and comments during the course of this project. We also thank FlyBase for maintaining databases and the Bloomington Drosophila Stock Center for providing fly stocks. The experiments were performed using the PiCSL-FBI core facility (IBDM, Marseille, France), a member of the France-BioImaging national research infrastructure supported by the French National Research Agency (ANR-10-INBS-04-01, “Investissements d’Avenir”). B.D. was supported by the ERC (grant Biomecamorph #323027) and Fondation Bettencourt Schueller. R.C. and J-M.P. were supported by the CNRS. TL was supported by the CNRS, followed by the Collège de France.

## Author contributions

B.D. and T.L. conceived the project. B.D. performed experiments/quantifications and developed analytical methods. R.C. designed the numerical model and performed the simulations. G.G-G. isolated the *frl*^59/59^ null allele. J-M.P. created all the fluorescent constructs. B.D., R.C. and T.L. discussed the data and wrote the manuscript.

---

**Supplementary Fig. 1.**
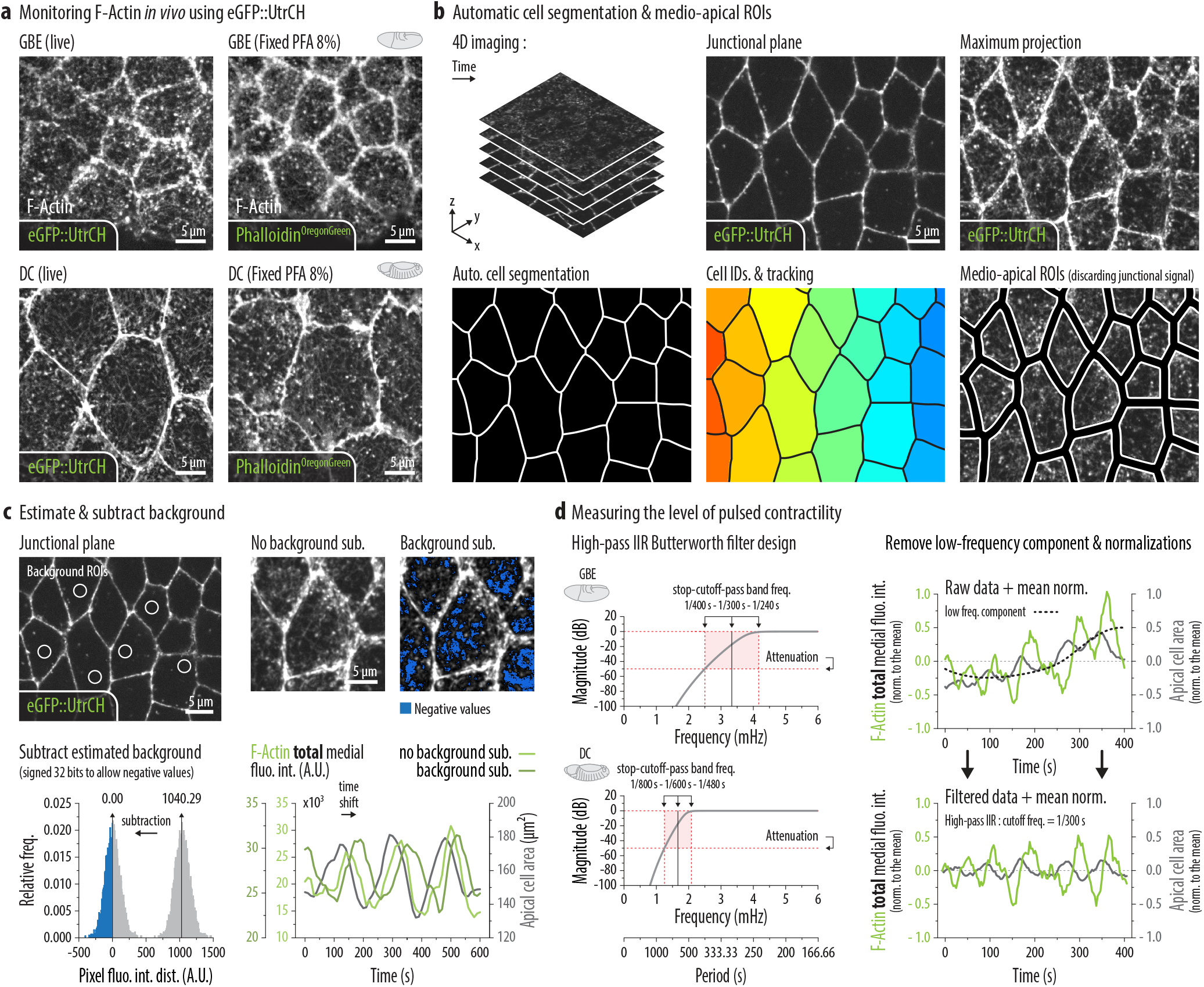
**(a)** Comparison between live (eGFP::UtrCH) and fixed (Phalloidin) F-Actin localization in ectodermal (GBE) and amnioserosa cells (DC). Images represent a max-proj. of 4 x 0.33 µm. **(b)** Presentation of the automatic cell segmentation procedure used to define medio-apical ROIs for quantification (see Methods for more details). Briefly, cell boundaries are detected on the lower junctional plane using a watershed algorithm. The segmented cells are then identified and tracked over time to define ROIs. Finally, these ROIs are shrunk of a few pixels to discard the junctional signal and the medial fluorescence intensities are measured on the max. proj. of the Z-series. **(c)** Presentation of the background subtraction procedure (see Methods for more details). The background is evaluated on the lower Z-planes and subtracted from the max. proj. of the Z-series before quantification. Removing the background is critical to properly measure the total amount of fluorescence in the ever-changing apical cell surface (see time shift when comparing the medial F-Actin levels with or without background subtraction). **(d)** To quantify the levels of pulsed contractility, we processed single cell profiles using a high-pass Butterworth IIR filter (see Methods for more details). This filter is used to remove low frequency components and have been adjusted to fit the temporality of pulsatility in GBE (cutoff freq. 1/300 s) and DC (cutoff freq. 1/600 s).

**Supplementary Fig. 2.**
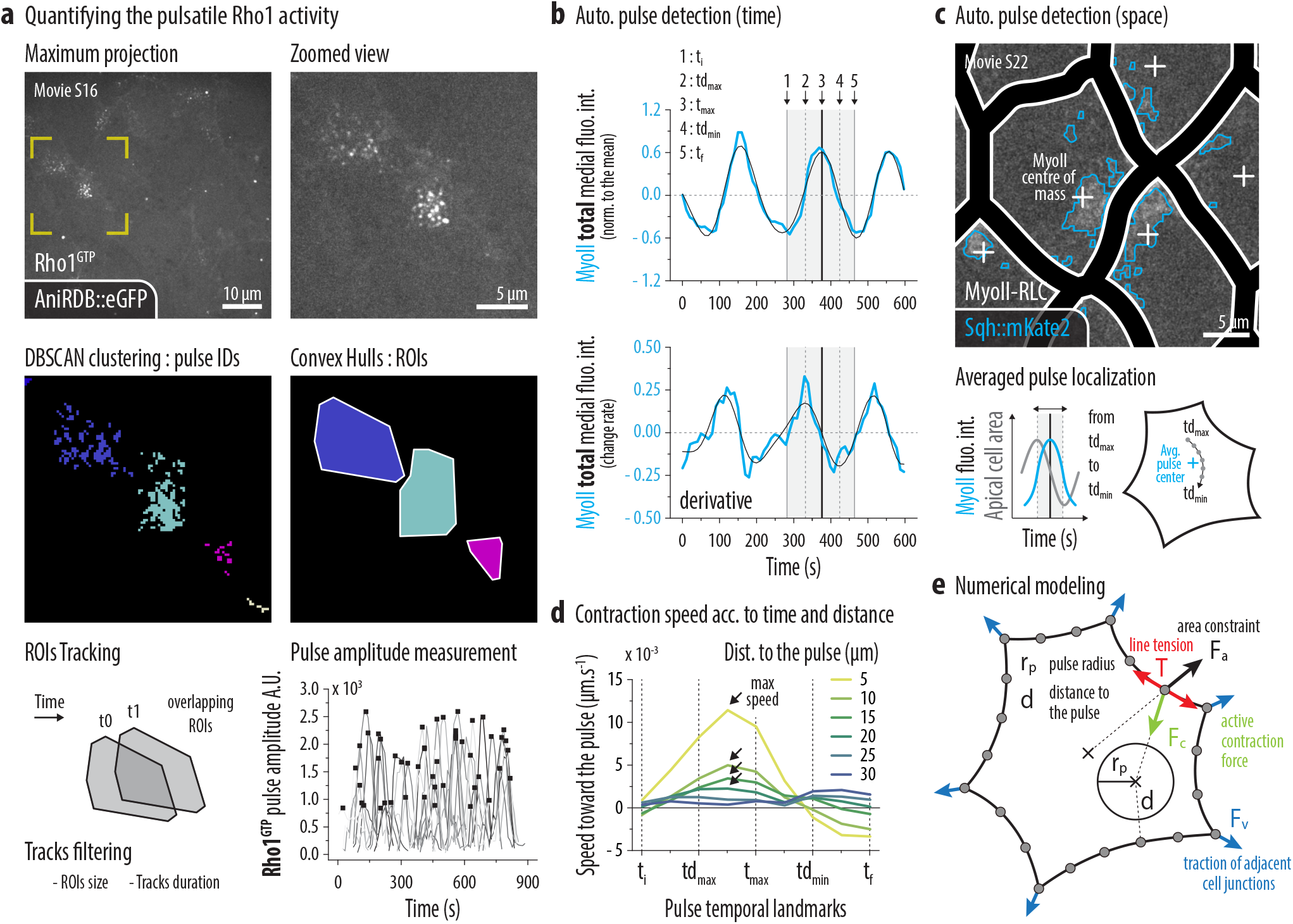
**(a)** Description of the method used to quantify to pulsatile Rho1 activity without cell segmentation (see Methods for more details and Supplementary Movie 16). Basically, isolated clusters of AniRBD::eGFP signal are detected using a DBSCAN algorithm. These clusters are then converted into surface ROIs using convex hulls and overlapping ROIs tracked over time to follow individual pulses. Filters such as min./max. area or min./max. duration are applied to reduce tracking mistakes. Finally, the AniRBD::eGFP pulse amplitude measurements are performed considering the maximum of total fluorescence intensity for each track. **(b)** MyoII pulses are automatically detected in time following the derivatives of high-pass filtered total medial MyoII levels (see Methods for more details). Pulse temporal landmarks have been defined as follow : ti : initial time; tdmax : max derivative; t_max_ : max amplitude; td_min_ : min derivative; t_f_ : final time. **(c)** MyoII pulses are automatically detected in space by monitoring the centre of mass of the medial sqh::mKate2 signal over time (see Methods for more details and Supplementary Movie 22). The actual pulse centre used for the KLT analysis is then defined by averaging the position of the recorded centre of mass between td_max_ and td_min_ of the pulse. This time interval corresponds to the period during which the apical surface contract during a pulse. **(d)** Line plots : Averaged speed towards the pulse according to pulse temporal landmarks (see above) for different distance bins (see legend). The black arrows show the time of maximum speed. **(e)** Schematics of the numerical model. The black circle depicts the actomyosin pulse, and arrows depict forces applied to boundary elements. Only a fraction of boundary elements is represented.

## Movie legends

**Supplementary Mov. 1 | F-Actin dynamics in ectodermal cells (GBE).** Live 100x imaging of F-Actin (eGFP::UtrCH) in ectodermal cells during germband extension (GBE). The movie represents a max. proj. of the 4 most apical z-planes, spaced by 0.33 µm and acquired every 3 seconds.

**Supplementary Mov. 2 | F-Actin dynamics in amnioserosa cells (DC).** Live 100x imaging of F-Actin (eGFP::UtrCH) in amnioserosa cells during dorsal closure (DC). The movie represents a max. proj. of the 4 most apical z-planes, spaced by 0.33 µm and acquired every 3 seconds.

**Supplementary Mov. 3 | Medial F-Actin turnover (GBE+DC).** High temporal resolution live 100x imaging of F-Actin (eGFP::UtrCH) in ectodermal cells during germband extension (GBE, left panel) and in amnioserosa cells during dorsal closure (DC, right panel). The movie represents a max. proj. of the 2 most apical z-planes, spaced by 0.33 µm and acquired every 1 second.

**Supplementary Mov. 4 | Medial MyoII and F-Actin dynamics (GBE).** Live 100x imaging of MyoII (Sqh::mCherry, left panel) and F-Actin (eGFP::UtrCH, right panel) in ectodermal cells during germband extension (GBE). The movie represents a max. proj. of the 4 most apical z-planes, spaced by 0.33 µm and acquired every 3 seconds.

**Supplementary Mov. 5 | Medial MyoII and F-Actin dynamics (DC).** Live 100x imaging of MyoII (Sqh::mCherry, left panel) and F-Actin (eGFP::UtrCH, right panel) in amnioserosa cells during dorsal closure (DC). The movie represents a max. proj. of the 4 most apical z-planes, spaced by 0.33 µm and acquired every 3 seconds.

**Supplementary Mov. 6 | Rho1 pathway inhibition (GBE).** Live 100x imaging of MyoII (Sqh::mCherry, top panels) and F-Actin (eGFP::UtrCH, bottom panels) in ectodermal cells during germband extension (GBE). Left panels : control embryo (water injected), middle panels : C3-transferase (Rho1 inhibitor) injected embryo, right panels : *RhoGEF2*^-/-^ embryo. The movies represent a max. proj. of the 4 most apical z-planes, spaced by 0.33 µm and acquired every 3 seconds.

**Supplementary Mov. 7 | Rho1 pathway inhibition (DC).** Live 100x imaging of F-Actin (eGFP::UtrCH) in amnioserosa cells during dorsal closure (DC). White outlines : control cells, yellow outline : Rho1N19 (Rho1 dominant negative form) expressing cell. The movie represents a max. proj. of the 4 most apical z-planes, spaced by 0.33 µm and acquired every 10 seconds.

**Supplementary Mov. 8 | Rok kinase inhibition (DC).** Live 100x imaging of MyoII (Sqh::mCherry, top panel) and F-Actin (eGFP::UtrCH, bottom panel) in amnioserosa cells during dorsal closure (DC). Left panels : control embryo (water injected), right panels : H-1152 (Rok inhibitor) injected embryo. The movies represent a max. proj. of the 4 most apical z-planes, spaced by 0.33 µm and acquired every 10 seconds.

**Supplementary Mov. 9 | Frl loss of function (DC).** Live 100x imaging of F-Actin (eGFP::UtrCH) in amnioserosa cells during dorsal closure (DC). Left panel : control embryo, right panel : Frl shRNA expressing embryo. The movie represents a max. proj. of the 4 most apical z-planes, spaced by 0.33 µm and acquired every 10 seconds.

**Supplementary Mov. 10 | Frl loss or gain of function (GBE).** Live 100x imaging of F-Actin (eGFP::UtrCH) in ectodermal cells during germband extension (GBE). Left panel : control embryo, middle panel : frl^59/59^ (null mutant) embryo, right panel : Frl^OE^ (overexpression) embryo. The movie represents a max. proj. of the 2 most apical z-planes, spaced by 0.33 µm and acquired every 2 seconds.

**Supplementary Mov. 11 | Frl loss or gain of function (DC).** Live 100x imaging of F-Actin (eGFP::UtrCH) in amnioserosa cells during dorsal closure (DC). Left panel : control embryo, middle panel : frl^59/59^ (null mutant) embryo, right panel : Frl^OE^ (overexpression) embryo. The movie represents a max. proj. of the 2 most apical z-planes, spaced by 0.33 µm and acquired every 2 seconds.

**Supplementary Mov. 12 | MyoII and F-Actin dynamics in Frl loss or gain of function (GBE).** Live 100x imaging of MyoII (Sqh::mKate2, top panel) and F-Actin (eGFP::UtrCH, bottom panel) in ectodermal cells during germband extension (GBE). Left panel : control embryo, middle panel : frl^59/59^ (null mutant) embryo, right panel : Frl^OE^ (overexpression) embryo. The movie represents a max. proj. of the 4 most apical z-planes, spaced by 0.33 µm and acquired every 6 seconds.

**Supplementary Mov. 13 | MyoII and F-Actin dynamics in Frl loss or gain of function (DC).** Live 100x imaging of MyoII (Sqh::mKate2, top panel) and F-Actin (eGFP::UtrCH, bottom panel) in amnioserosa cells during dorsal closure (DC). Left panel : control embryo, middle panel : frl^59/59^ (null mutant) embryo, right panel : Frl^OE^ (overexpression) embryo. The movie represents a max. proj. of the 4 most apical z-planes, spaced by 0.33 µm and acquired every 10 seconds.

**Supplementary Mov. 14 | Frl gain of function (DC).** Live 100x imaging of F-Actin (eGFP::UtrCH) in amnioserosa cells during dorsal closure (DC). White outlines : control cells, yellow outline : Frl^OE^ (overexpression) cell. The movie represents a max. proj. of the 4 most apical z-planes, spaced by 0.33 µm and acquired every 10 seconds.

**Supplementary Mov. 15 | Rho1GTP dynamics in Frl loss or gain of function (DC).** Live 100x imaging of Rho1GTP (AniRBD::eGFP) in amnioserosa cells during dorsal closure (DC). Left panel : control embryo, middle panel : Frl shRNA expressing embryo, right panel : Frl^OE^ (overexpression) embryo. The movie represents a max. proj. of the 4 most apical z-planes, spaced by 0.33 µm and acquired every 10 seconds.

**Supplementary Mov. 16 | Automatic Rho1GTP pulse tracking (DC).** Live 100x imaging of Rho1GTP (AniRBD::eGFP) in amnioserosa cells during dorsal closure (DC), showing the method used to automatically track Rho1GTP pulses without cell segmentation. Left panel : tracked ROIs, right panel : individual pulses detected using DBScan clustering. The movie represents a max. proj. of the 4 most apical z-planes, spaced by 0.33 µm and acquired every 10 seconds.

**Supplementary Mov. 17 | Epithelial dynamics in Frl loss or gain of function (GBE).** Live 40x imaging of F-Actin (eGFP::UtrCH) in ectodermal cells during germband extension (GBE). Left panel : control embryo, middle panel : frl^59/59^ (null mutant) embryo, right panel : Frl^OE^ (overexpression) embryo. The yellow cell outlines represent the results of cell segmentation and the white squares mark the localization of T1 events. The movie represents one z-plane, acquired every 20 seconds.

**Supplementary Mov. 18 | Germband extension in Frl loss or gain of function (GBE).** DIC live imaging of embryos undergoing germband extension (GBE). Top panel : control embryo, middle panel : frl^59/59^ (null mutant) embryo, bottom panel : Frl^OE^ (overexpression) embryo. The movie represents one z-plane, acquired every 30 seconds.

**Supplementary Mov. 19 | Apical cell surface deformations in Frl loss or gain of function (DC).** Live 100x imaging of F-Actin (eGFP::UtrCH) in amnioserosa cells during dorsal closure (DC). Left panel : control embryo, middle panel : frl^59/59^ (null mutant) embryo, right panel : Frl^OE^ (overexpression) embryo. The inserted images represent the results of cell segmentation. The movie represents a max. proj. of the 4 most apical z-planes, spaced by 0.33 µm and acquired every 10 seconds.

**Supplementary Mov. 20 | Dorsal closure in Frl loss or gain of function (DC).** Live 10x imaging of F-Actin (eGFP::UtrCH) of embryos undergoing dorsal closure (DC). Top panel : control embryo, bottom panel : Frl shRNA expressing embryo. The movie represents a max. proj. of the 10 z-planes, spaced by 5 µm and acquired every 10 minutes.

**Supplementary Mov. 21 | Contractile event dynamics in Frl loss of function (DC).** Live 100x imaging of F-Actin (eGFP::UtrCH) in amnioserosa cells during dorsal closure (DC). Left panel : control embryo, right panel : frl^59/59^ (null mutant) embryo. The movie represents a max. proj. of the 2 most apical z-planes, spaced by 0.33 µm and acquired every 2 seconds.

**Supplementary Mov. 22 | Automated pulse and KLT tracking (DC).** Live 100x imaging of MyoII (Sqh::mKate2, left panel) and F-Actin (eGFP::UtrCH, right panel) in amnioserosa cells during dorsal closure (DC). Left panel : automated MyoII pulse tracking in space, the white crosses represent the medial MyoII centre of mass. Right panel : F-Actin KLT tracked particles, the color code represents the speed of tracked particles in µm.s^-1^. The movie represents a max. proj. of the 4 most apical z-planes, spaced by 0.33 µm and acquired every 5 seconds.

**Supplementary Mov. 23 | Numerical model.** Representative simulations for different values of *λ* (10 examples per condition). The pulse is represented by the inner circle and the green segments indicate that a boundary element is connected to the pulse. The pulse position is chosen randomly in the different examples.

## Materials and Methods

### Fly strains and genetics

We visualized the F-Actin dynamics in living embryos using a sqh-eGFP::UtrCH (Calponin Homology domain of Utrophin) insertion either on the 2^nd^ or 3^rd^ chromosome^7^. To co-image the F-Actin with the MyoII, we recombined the eGFP::UtrCH with a fluorescent version of the *Drosophila* MyoII-RLC (encoded by the *spaghetti-squash* or *sqh* gene) using a sqh-Sqh::mCherry or a sqh-Sqh::mKate2 insertion. In both cases, these Sqh constructs were inserted either on the K18 site (53B2) on the 2^nd^ chromosome or the VK27 (89E11) on the 3^rd^ chromosome. To monitor the Rho1 GTPase activity *in vivo* we used the Rho1GTP sensor ubi-AnillinRBD::eGFP (Rho Binding Domain of anillin) inserted on the 3^rd^ chromosome^13^.

The 67-Gal4 (mat-4-GAL-VP16) or the *engrailed*-GAL4 (en2.4-GAL4e16E, UAS-NLS::RFP, BDSC #30557) drivers, carried on the 2^nd^ chromosome, have been combined to sqh-eGFP::UtrCH, sqh-Sqh::mKate2 or ubi-AnillinRBD::eGFP on the 3^rd^ chromosome and were used to express the following constructs : UAS-Rho1N19 (BDSC #58818), UAS-Frl^WT^ (gift from Andreas Jenny)^45^ and UAS-Frl^shRNA^ (CG32138 TRiP line, BDSC #32447). The RhoGEF2 germline clones, using the RhoGEF2^l^(2)^04291^ null allele^54^, have been made using the FLP-DFS system^55^.

To visualize the effect of a ubiquitous overexpression of Frl (Frl^OE^) in ectodermal cells during GBE we crossed 67-GAL4/+; UAS-Frl^WT^/+ females with UAS-Frl^WT^/UAS-Frl^WT^ males (maternal/zygotic Frl overexpression). To visualize the effect of a ubiquitous overexpression of Frl (Frl^OE^) in amnioserosa cells during DC we crossed 67-GAL4/67-GAL4; +/+ females with UAS-Frl^WT^/UAS-Frl^WT^ males (zygotic Frl overexpression). To reduce the endogenous Frl levels using shRNA we crossed 67-GAL4/+; UAS-Frl^shRNA^/+ females with UAS-Frl UAS-Frl^shRNA^/UAS-Frl UAS-Frl^shRNA^ males (maternal/zygotic Frl shRNA expression).To drive the expression of Rho1N19 or the overexpression of Frl in isolated amnioserosa cells during DC, we respectively crossed *engrailed*-GAL4, UAS-NLS::RFP/CyO; +/+ females with UAS-Rho1N19/CyO or UAS-Frl^WT^/UAS-Frl^WT^ males (zygotic mosaic expression/overexpression). The nuclear NLS::RFP signal have been used as a reporter to identify the isolated amnioserosa cells expressing the *engrailed*-GAL4 driver. Looking at the distribution of nuclear NLS::RFP intensities we defined a threshold to categorize cells between controls and overexpressing cells (see Fig. 3b).

### Constructs and transgenesis

The *frl*^59^ mutant was generated by the CRISPR/Cas9 technique^56^. In brief, two 21 nt long gRNAs, GAGCAACTTTGCTTTATCCGG and GTCGTTTATCGCGCACCCTGG, were designed with homology to the second and last coding exons of frl, respectively, and cloned into the pCFD4 vector. After germ cell-specific simultaneous expression of Cas9 and the gRNAs, we collected frl mutant candidates from the second generation which were validated by PCR and sequencing. Based on the sequencing data, the expected ∼ 8640 bps deletion was detected from the genomic DNA of the mutant strains. We next associated the null *frl*^59^ allele (3^rd^ chromosome) with a sqh-eGFP::UtrCH, sqh-Sqh::mKate2 recombinant carried on the 2^nd^ chromosome. From this stock, we selected male and female adult fly homozygote for Frl^59^ and cross them together to study the effect of a maternal/zygotic depletion of Frl on the actomyosin dynamics.

### Live imaging

Embryos were prepared for live imaging as previously described^57^ and movies were acquired at room temperature (22°C) at stage 7-8 for ectodermal cells during GBE and at stage 13-14 for amnioserosa cells during DC. All live imaging has been performed using a dual camera (QImaging, Rolera EM-C^2^, EMCCD) spinning disc (CSU-X1, Yokogawa) on a Nikon Eclipse Ti inverted microscope (distributed by Roper) managed by the MetaMorph software. Dual color imaging of eGFP and mCherry/mKate2 FPs was obtained by simultaneously exciting fluorophores with a 491 nm and a 561 nm laser and using a dichroic mirror to collect emission signals on two cameras. We used the following Nikon objectives : 10X/N.A. 0.25 dry, 40X/N.A. 1.25 water and a 100X/N.A. 1.4 oil. For the 100X movies, we focused on the most apical part of epithelial cells, performing Z-series of 1 to 6 planes separated by 0.33 µm and acquired every 1 to 10 seconds (see exact imaging conditions in the supplementary movies legend). For the 40X movies (Fig. 6c-f), we searched to monitor epithelial dynamics and filmed for that only one optical section at the AJs level every 20 seconds. For the 10X movies (Fig. 6g-i), we imaged Z-series of 10 planes separated by 5 µm every 3 minutes, to capture most of the embryo volume over the duration of the DC process. In all cases, imaging conditions (exposure time, laser power) were optimized and kept constant between controls and perturbed embryos.

### Drug injections

To inhibit the Rho1 GTPase in ectodermal cells during GBE, we injected the C3-transferase exoenzyme (from Cytoskeleton, Inc), resuspended in water at 0.5 µg/µl, in the yolk of pre-gastrulating embryos. Injections had to be performed just before the end of cellularization to allow this non-cell permeable drug to penetrate the cells without impairing, too early, the first movement of gastrulation. To inhibit the Rok kinase in amnioserosa cells during DC, we injected the H-1152 compound (from Tocris Bioscience), resuspended in water at 40 mM, in the perivitelline space of embryos. Before the injections, the embryos were slightly dried by being exposed during ~ 7 min to the Drierite (Sigma-Aldrich), to prevent cells from being expelled from the vitelline shell. Injections have been performed on the imaging microscope using an InjectMan4 micromanipulator and a FemtoJet 4i microinjector from Eppendorf. Embryos were imaged either ~ 10 min (C3-transferase) or ~ 2 min (H-1152) after injection.

### Embryos fixation and phalloidin staining

To validate the eGFP::UtrCH probe, we compare the live localization of F-Actin obtained using the probe with fixed embryos stained by phalloidin (Supplementary Fig. 1a). To do so, we fixed embryos in a half-half mix of heptane and 8% paraformaldehyde (diluted in PBS) for 30 minutes under constant shaking. The embryos were then washed in PBS 10% BSA and hand devitellinized using a thin syringe needle. We next incubated embryos for 2 hours at room temperature in a blocking/permeabilizing PBS 10% BSA + 0.3% triton X-100 solution. The OregonGreen 488 phalloidin (from Invitrogen) was diluted 1/50 in PBS 10% BSA and the staining was performed at room temperature for 30 minutes. Before mounting, the embryos were washed a last time in PBS without BSA. Imaging was performed on a Leica LSM SP8 microscope using a 100X/N.A. 1.4 oil objective.

### Image processing and data analysis

#### Used software

All image processing and data analysis have been performed using the ImageJ or Matlab software, either separately or together using the MIJ plugin (D.Sage, D.Prodanov, C.Ortiz and JY.Tivenez, retrieved from http://bigwww.epfl.ch/sage/soft/mij). The graphics were produced using OriginPro software and exported to Adobe Illustrator for final processing.

#### Automatic cell segmentation

To measure fluorescence intensities in the medio-apical cortex, we first designed an automated cell segmentation procedure based on watershed algorithms (Supplementary Fig. 1b). To achieve this, we used the eGFP::UtrCH signal from the lower planes of our Z-series (AJs planes) and reduce the images size (x 0.33) to speed up the procedure. We then run the DIPimage watershed algorithm (retrieved from http://www.diplib.org/dipimage) and made a custom Matlab/ImageJ code to select and track segmented cells over time. If required (especially for the 40x movies), we manually corrected the segmentation results using the Tissue Analyzer ImageJ plugin (B.Aigouy, retrieved from https://grr.gred-clermont.fr/labmirouse/software/WebPA)^58^. The Tissue Analyzer plugin was also used in Fig. 6c-f for automatic detection of irreversible T1 events (cell intercalation).

#### Image processing and measurements

To prepare the 100X images for quantification we first maximum-projected our Z-series over the 2 to 4 most apical planes. We then evaluated and subtracted the cytoplasmic background by measuring fluorescence intensities on the lower Z-planes. For this purpose, we projected the lower planes as we did for the apical planes and manually drew circular ROIs to measure the cytoplasmic levels (Supplementary Fig. 1c). This step is critical to properly measure the total amount of fluorescence in the ever-changing apical cell surface. For example, without background subtraction, pulses cannot be properly timed since the fluctuations of the apical surface introduce a shift in the measurement of medial fluorescence levels. Finally, we measured the medio-apical fluorescence intensities using the ROIs obtained by segmenting the cells and shrunk these ROIs, by 10 pixels for ectodermal cells or 15 pixels for amnioserosa cells, to prevent the junctional signal from contaminating the measurements. According to cases, we either measured the mean or the integrated medial fluorescence intensities to, respectively, get information about the density or the total levels of the considered fluorescent protein.

#### Measuring the level of pulsed contractility

During this study we repeatedly assessed the level of pulsed contractility by comparing the medial actomyosin pulsatility and the apical cell area fluctuations (Fig. 2f; 3d,g; 5c,d,g; 6e). To do so, we measured the standard deviations (SDs) of individual cell profiles after normalizing data to the mean and removing the low frequency components using an IIR Butterworth high-pass filter (Matlab signal processing toolbox). We choose a cutoff frequency (1/300s for ectodermal cells during GBE and 1/600s for amnioserosa cells during DC) that we multiply by 0.75 or 1.25 to respectively define the stop and the pass band frequency (Supplementary Fig. 1d). The high-pass filter allowed us to isolate the effect of pulsed contractility by discarding the contribution of slower variations of the apical surface. Indeed, it appeared that the apical cortex tends to steadily increase/decrease its surface over the time of recording (~ 450 seconds for GBE or ~ 900 seconds for DC) influencing, therefore, our SDs measurements. Removing the low frequency components revealed particularly required to measure the F-Actin pulsatility since the density of medial filaments tend to scale with the apical cell surface.

#### Cross-correlation analysis

To evaluate how the pulsatile F-Actin accumulations synchronizes with the MyoII pulses and the cycles of apical constriction we performed a cross-correlation analysis, comparing the variation of the total medial F-Actin levels with either the variation of the total medial MyoII levels or the fluctuation of apical cell area (Fig. 1i,j). In both cases, we pre-processed our data using a mean normalization and a high-pass filter (see above). The normalized cross-correlation have been calculated for individual cells using the following formula, where *t* is the time, *T* is the total time of analysis and *τ* is the time delay:

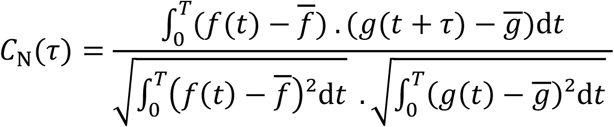

The mean cross-correlation curves have been obtained by averaging the individual cell results and the typical time delay have been plotted considering the maximum correlation for the F-Actin/MyoII comparison and the minimum correlation for the F-Actin/apical cell area comparison.

#### Quantifying the pulsatile Rho1 activity

To quantify the levels of Rho1 activity during pulses we measured the amplitude of AniRBD::eGFP accumulations during contractile events (Rho1GTP sensor) (Fig. 5h-j). However, since we were unable to segment cells using the AniRBD::eGFP signal, we designed a live pulse tracking procedure working directly on unsegmented images (Supplementary Fig. ??). To do so, we used a Matlab implementation of the DBSCAN clustering algorithm (S. Mostapha Kalami Heris, retrieved from http://yarpiz.com/255/ypml110-dbscan-clustering) integrated to a custom code for automatization. This method allowed us to isolate clusters of AniRBD::eGFP signal based on two main parameters : the distance ε and the minimum number of points that must be within an ε radius for these points to be considered as a cluster. We then converted these clusters into ROIs, using convex hulls (minimum area polygon containing the cluster), and tracked overlapping ROIs over time to follow individual pulses. After filtering pulses, based on ROIs sizes and tracks duration criterions, we measured the amplitude of AniRBD::eGFP pulses by measuring the maximum of total fluorescence intensity for each track.

#### Cell shape regularity measurements

To measure how regularly cells were shaped, we compared the segmented cell boundary with the convex hull formed by connecting vertices (Fig. 6j-l; 7g). We next quantified the area occupied by the inward/outward convolutions of the actual cell shape and divided it by the surface of the convex hull. In doing so, we obtained a ratio whose value indicates the convolution level of the segmented cell shape (the higher the value the more convoluted is the shape). We calculated this ratio for every cell at every time point and then compare it between conditions.

#### KLT / pulse analysis

To quantify the propagation of MyoII pulsatile stresses within the cortex, we measured the speed at which tracked apical F-Actin structures (eGFP::UtrCH signal) displace toward the pulse center during contractile events. We first tracked these apical F-Actin structures using a Kanade-Lucas-Tomasi features tracking algorithm (KLT)^59, 60^ implemented in C (S.Birchfield, retrieved from https://cecas.clemson.edu/~stb/klt) whose tracking results have been exported to Matlab for further processing. Briefly, the KLT algorithm first detects discrete features within the image by examining the minimum eigenvalue of each 2 by 2 gradient matrix. These features are then tracked over time following a Newton-Raphson method of minimizing the difference between the two windows. Finally, to minimize tracking mistakes, each feature is checked by an affine consistency test, comparing the spatial distribution of the feature and its neighbors at given time-point to the time-point at which the feature was first detected^61^. In our case, since the medio-apical F-Actin is in a constant reorganization, we were not able to follow features over long periods and therefore had to replace them during the movie in order to produce enough data. Considering only the tracks whose duration exceed three time-points, we obtained in average ~ 50,000 tracks per embryo, each of these containing the x y coordinates of the tracked feature.

We next sought to detect pulses, in time and space, in order to define the reference points from which our propagation measurements were made. To do so, we first designed an automated pulse detection procedure, working on the smoothed derivatives of high-pass filtered total medial MyoII levels (see Auto. pulse detection (time) in Supplementary Fig. 2b). This procedure, based on the “findpeaks” function of the Matlab Signal Processing Toolbox, allowed us to detect and define temporal landmarks within pulses (t_i_ : initial time; td_max_ : max derivative; t_max_ : max amplitude; td_min_ : min derivative; t_f_ : final time) and to adjust the detection sensitivity using parameters such as the peak prominence, the peak width and the peak prominence over width ratio. In a second time, we implemented another Matlab routine to monitor the medial MyoII center of mass within contracting cells (see Auto. pulse detection (space) in Supplementary Fig. 2c). To perform this analysis, we first removed the junctional MyoII (post-processing) and used a combination of gaussian blurs and thresholding steps to isolate the MyoII signal actually belonging to a pulse. Once defined, we averaged the positions of the MyoII center of mass between the t_dmax_ and td_min_ time-points and used this reference point (pulse center) for our propagation measurements (Fig. 7d).

Finally, to produce the propagation curves in Fig. 7e we averaged the speed, between tdmax and td_min_, at which the KLT tracked apical F-Actin structures converge toward the pulse center (define as describe above). This averaging step allowed us to smooth the curve profile and to focus only on the period corresponding to the apical cell contraction (convergence of apical F-Actin structures). We then binned data according to the distance and plotted the measured speed toward the pulse as a function of the distance to the pulse.

### Numerical modeling

We consider a single cell submitted to a contraction due to an actomyosin pulse. The cell is represented by a discretized contour of N attachment sites (we use N=100 in the simulations). Six of the sites represent the vertices of the cell. Initially, they are regularly distributed along the contour. The sites are submitted to the following forces:

- An <u>area constraint</u> in the form of cell area elasticity:

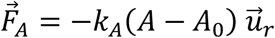

*k_A_* is the area stiffness, i.e. the strength associated to the area constraint, *A* is the area of the cell, *A*_0_ its target area, and 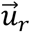 the unit radial vector. The origin is taken at the cell center of mass.

- A <u>line tension</u> along the cell contour. The site *i* pulls on the neighboring sites *i* − 1 and *i* + 1:

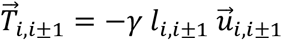

*γ* is the line tension constant, *l*_*i, i* ± 1_ the distance between sites *i* and *i* ± 1, and 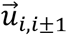 the unit vector along the segment joining sites *i* and *i* ± 1.

- To mimic the <u>traction of adjacent cell junctions</u> on vertices, we assume that sites corresponding to vertices undergo an additional elastic force:

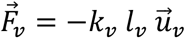

*k_v_* is the strength associated to the elastic force, and *l_v_* the distance of the vertex to its initial position. 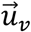 is the unit vector along the segment joining the vertex to its initial position.

- An <u>active contraction force</u> distributed among the connected attachment sites. The contraction force on site *i* is:

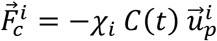

Where *χ*_i_ is either 1 or 0 (connected / not connected, see below), *C*(*t*) is the amplitude of the contraction force, and 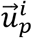 is the unit vector along the segment joining site *i* to the pulse center.

In each simulation, the position of the pulse center is chosen randomly anywhere inside the cell. The probability that a site of the cell contour is connected to the pulse depends on its distance *d_i_* to the pulse:

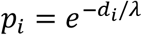

*λ* is a connectivity length scale, and a direct indicator of network density. A sparse network will result in a short *λ*, while a dense network will result in a long *λ*. Note that we consider that the pulse has a non-zero spatial extension, with radius *r_p_*. The probability that an attachment site within the pulse radius is connected to the pulse is equal to 1. Hence the distance *d_i_* is computed as the distance to the pulse circumference.

The amplitude of the contraction force *C*(*t*) is:

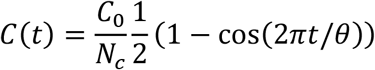

The amplitude on each site *C_0_* /*N_c_* is normalized to the number of connected sites *N_c_* so that the total work produced does not depend on the number of connected sites. We assume that the contraction amplitude oscillates with period *θ*. Due to the periodicity, we limit the simulations to one period.

The dynamics is simulated by solving the force balance equation, assuming a fluid friction force with friction coefficient *α*:

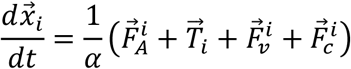

In our simulations, we used *A*_0_ = 170 μm^3^, *r_p_* = 3.5 μm, *θ* = 200 s, *γ* = 120, *α* = 5, *k_A_* = 0.05, *k_ν_* = 30 and *C*_0_ = 500. Note that with these parameters, the initial (equilibrium) configuration before the contraction starts is a hexagon. We used different values of *λ* ranging from 2*μm* to 100*μm* to illustrate the role of connectivity.

### Statistics

Data points from different pulses/cells/embryos were pooled to estimate the mean, S.D. and S.E.M. Statistical significances have been tested using two-sample t-test, assuming normal distributions and une qual variances. No statistical method was used to predetermine sample size. The experiments were not randomized, and the investigators were not blinded to allocation during experiments and outcome assessment.

### Data and codes availability

The authors declare that the data and the analysis methods supporting the findings of this study are available within the paper and its supplementary information files. Raw image data and code are available upon request.

